# Sex-Specific Effects of Early-Life Iron Deficiency and Prenatal Choline Treatment on Adult Rat Hippocampal Transcriptome

**DOI:** 10.1101/2022.06.29.498159

**Authors:** Shirelle X. Liu, Tenille K. Fredrickson, Natalia Calixto Mancipe, Michael K. Georgieff, Phu V. Tran

## Abstract

**Background:** Fetal-neonatal iron deficiency (ID) causes long-term neurocognitive and affective dysfunctions. Clinical and preclinical studies have shown that early-life ID produces sex-specific effects. However, little is known about the molecular mechanisms underlying these early-life ID-induced sex-specific effects on neural gene regulation.

**Objective:** To illustrate sex-specific transcriptome alteration in adult rat hippocampus induced by fetal-neonatal ID and prenatal choline treatment.

**Methods:** Pregnant rats were fed an iron-deficient (4 mg/kg Fe) or iron-sufficient (200 mg/kg Fe) diet from gestational day (G) 2 to postnatal day (P) 7 with or without choline supplementation (5 g/kg choline) from G11-18. Hippocampi were collected from P65 offspring of both sexes and analyzed for changes in gene expression.

**Results:** Both early-life ID and choline treatment induced transcriptional changes in adult female and male rat hippocampus. Both sexes showed ID-induced alterations in gene networks leading to enhanced neuroinflammation. In females, ID-induced changes indicating enhanced activity of oxidative phosphorylation and fatty acid metabolism, which are contrary to the ID effect in males. Prenatal choline supplementation induced the most robust changes in gene expression, particularly in the iron-deficient animals where it partially rescued ID-induced dysregulations. Choline supplementation also altered hippocampal transcriptome in the iron-sufficient rats with indications for both beneficial and adverse effects.

**Conclusions:** This study provided unbiased global assessments of gene expression regulated by iron and choline status in a sex-specific manner, with greater effects in female than male rats. Our new findings highlight potential sex-specific gene networks regulated by iron and choline status for further investigation.

## INTRODUCTION

Iron deficiency (ID) during the fetal-neonatal period can result in long-term neurocognitive and emotional abnormalities [1–4]. In clinical populations, there is evidence that supports sex-specific outcomes of early-life ID and its treatments, including differential effects on fecal microbiota [5], brain development [6], verbal fluency [7], and risks of cognitive disorders such as schizophrenia [2]. Sex-specific physiologic and behavioral effects are also observed in preclinical models [8–11]. Early postnatal ID anemia causes sex-specific neuroinflammatory response [8], seizure vulnerability [9], social and affective behavioral abnormalities [10], and vasculature dysfunction [11]. However, analysis of sex-specific changes in the regulation of gene transcription that might underlie the neurodevelopmental effects of ID remains limited to a report of alterations in the hippocampal transcriptome from our phlebotomy-induced ID anemia (PIA) mouse model [8] and in the placental transcriptome from a maternal ID mouse model [12]. Thus, additional unbiased global transcriptomic studies are needed to establish the sex-specific molecular changes induced by early-life ID that may explicate the differential long-term neurobehavioral effects and disease vulnerabilities between sexes [13].

Hippocampus is an important brain structure that mediates learning, memory, and cognitive functions, which can be influenced by sex [14, 15]. The concurrent effects of early-life ID on the development and function of the rat hippocampus have been well documented [16–21]. However, long-term changes in transcriptional regulation induced by early-life ID between sexes within the hippocampus in rat models remain understudied, hindering a comprehensive understanding of the negative and long-lasting effects of early-life ID, as well as the development of more effective therapeutics.

Transcriptomic analysis using the Next-Generation Sequencing (NGS) technology offers a unique unbiased approach to identify global changes in the transcriptome. NGS findings can be annotated into changes in relevant biological processes, canonical pathways, and disease states utilizing the knowledge-based databases such as Ingenuity Pathway Analysis (IPA, Qiagen Inc.), Molecular Signatures Database (MSigDB), and Gene Ontology (GO) [22–24]. The present study analyzed and compared long-term changes in hippocampal gene regulation between adult male and female rats that were iron-deficient versus iron-sufficient during fetal-neonatal period and between those with and without prenatal choline treatment.

## MATERIALS AND METHODS

### Animals

Gestational day (G) 2 pregnant Sprague–Dawley rats were purchased from Charles River Laboratories (Wilmington, MA, USA). Rats were maintained in a 12-hr:12-hr light/dark cycle with ad lib food and water. ID was induced by dietary manipulation as described previously [21]. In brief, to generate iron-deficient pups, pregnant dams were given a purified iron-deficient diet (4 mg Fe/kg, TD.80396; Harlan Teklad) from G2 to postnatal day (P) 7 when lactating dams were switched to a purified iron-sufficient diet (200 mg Fe/kg, TD.09256; Harlan Teklad). Control iron-sufficient pups were generated from pregnant dams maintained on the iron-sufficient diet. Both diets were the same in all respects except the iron (ferric citrate) content. Half of the dams on the iron-sufficient or iron-deficient diet received dietary choline supplementation (5.0 g/kg choline chloride supplemented; iron-sufficient with choline: TD.1448261, iron-deficient with choline: TD.110139; Harlan Teklad) from G11-G18, while the other half of the dams received their iron-sufficient or iron-deficient diet with standard choline content (1.1 g/kg). Thus, dams and their litters were randomly assigned to one of the four groups based on the maternal diet: iron-deficient with choline supplementation (IDch), iron-deficient without supplemental choline (ID), always iron-sufficient with choline supplementation (ISch), always iron-sufficient without supplemental choline (IS). Details of the diet contents have been described in our previous study [25]. All litters were culled to 8 pups with equal number of each sex at birth. Rats were weaned at P21. Littermates were separated by sex and housed in groups of two. To avoid litter-specific effects, two rats per litters and ≥ 3 litters/group were used in the experiments. The University of Minnesota Institutional Animal Care and Use Committee approved all experiments in this study (Protocol # 2001-37802A).

### Hippocampal dissection

Rats were euthanized on P65 by an injection of pentobarbital (100 mg/kg, intraperitoneal). Brains were removed and bisected along the midline on an ice-cold metal block. Hippocampus was dissected and immediately flash-frozen in liquid nitrogen, banked and stored at −80°C for further use.

### RNA isolation and sequencing

Total RNA was isolated from the hippocampus of P65 females of all four groups (n=4/group) using the RNeasy Midi Kit (Qiagen). RNA samples were submitted to the University of Minnesota Genomics Center for library preparation and sequencing. RNA quantity and quality were assessed using the RiboGreen RNA Assay kit (Invitrogen) and capillary electrophoresis (Agilent BioAnalyzer 2100, Agilent), respectively. RNA samples with a RIN score > 8.0 were used for library preparation. Barcoded libraries were constructed for each sample using the TruSeq RNA v2 kit (Illumina). Libraries were size-selected for ~ 200bp fragments. Sequencing was performed using Illumina NovaSeq to generate 150bp pair-end reads. Sequencing depth was >25 million reads per sample for all samples.

### Bioinformatics

NGS reads from the P65 male hippocampi in a prior study [26] were sequenced with 50bp pair-end reads on a HiSeq2000 instrument. We hypothesize that changes in gene expression due to maternal ID and choline supplementation would have both sex-dependent and -independent responses. The sex-specific responses would contribute to differential outcomes of ID and choline supplementation between sexes. Because the data were collected in two batches with different sequencing technologies, the sex-diet interaction effects could be confounded. The following data preprocessing and analyses were performed: (1) Filtering and normalization - raw read counts were filtered to include only genes that were 300 bp or longer and expressed in at least one of the four treatment groups; (2) Statistical modeling – for differential gene expression tests, expression values were modeled by a generalized linear model (GLM) with a negative binomial distribution using either the dietary group as the explanatory variable if the model was built for each sex separately (16 RNA-sequencing (RNA-seq) libraries), or with the factors (iron, choline, sex, and their interactions) as explanatory variables, using all 32 RNA-seq libraries. For the analysis of variance, a linear mixed model with the factors (sex, choline, iron, and their interactions) was used; (3) Analysis of differentially expressed (DE) genes – differential gene expression was tested using a quasi-likelihood F-test on the fitted GLMs and the following comparisons: (ID, IDch) vs (IS, ISch) for the general iron effects, choline-supplemented (ISch, IDch) vs non-supplemented (IS, ID) for the general choline effects, iron by choline interaction, iron by sex interaction (sex-specific iron effects), and choline by sex interaction (sex-specific choline effects); genes with FDR < 0.05 were taken as DE between comparison groups. These genes were used to perform an unsupervised hierarchical clustering of the samples and a heatmap depicting the counts per million (CPM) values of the DE genes in each group. (4) Analysis of variance - the contributions of each factor to gene variances were estimated using the R package variancePartition (VP). The top 2% genes (approximately 300 genes) with variances explained primordially by the study variables (i.e., iron, choline, or sex) were selected for overrepresentation analysis (ORA) of GO terms. (5) Gene set enrichment analysis (GSEA) – all expressed genes were ranked using their differential expression test results by -log(p-value) * sign(fold change). Ranked gene lists were tested for enrichment against the Hallmark gene sets from MSigDB. An FDR < 0.25 cutoff was used for statistically significant set enrichment. A complete description of analyses and results is shown in supplementary materials (Supplemental Method).

### Ingenuity Pathway Analysis (IPA)

DE genes and ranked gene lists used in GSEA were mapped onto known biological processes using the “Core Analysis” feature of IPA, which includes > 8.5 million studies (Qiagen, Redwood City, CA). Ranked gene lists were also used to identify altered upstream regulators and known biomarkers in the hippocampus, cerebrospinal fluid (CSF), blood, and plasma/serum. Significant findings were determined using an absolute z-score ≥ 2.0 computed by Fisher’s exact test.

## Results

### Maternal ID and choline supplementation altered offspring’s hippocampal gene expression in a sex-specific manner

Effects of maternal ID: NGS data from both sexes were combined to assess the long-term effects of maternal ID on the transcriptional regulation of the offspring hippocampus in adulthood. After filtering and normalization, the combined dataset contained 15,060 loci. When the combined dataset was analyzed by a GLM to test for DE genes between iron-deficient (ID, IDch, both sexes) vs iron-sufficient (IS, ISch, both sexes) groups, gene expression changes failed to reach the FDR threshold (<0.05) in response to maternal iron status (Figure 1A).

**Figure 1:**
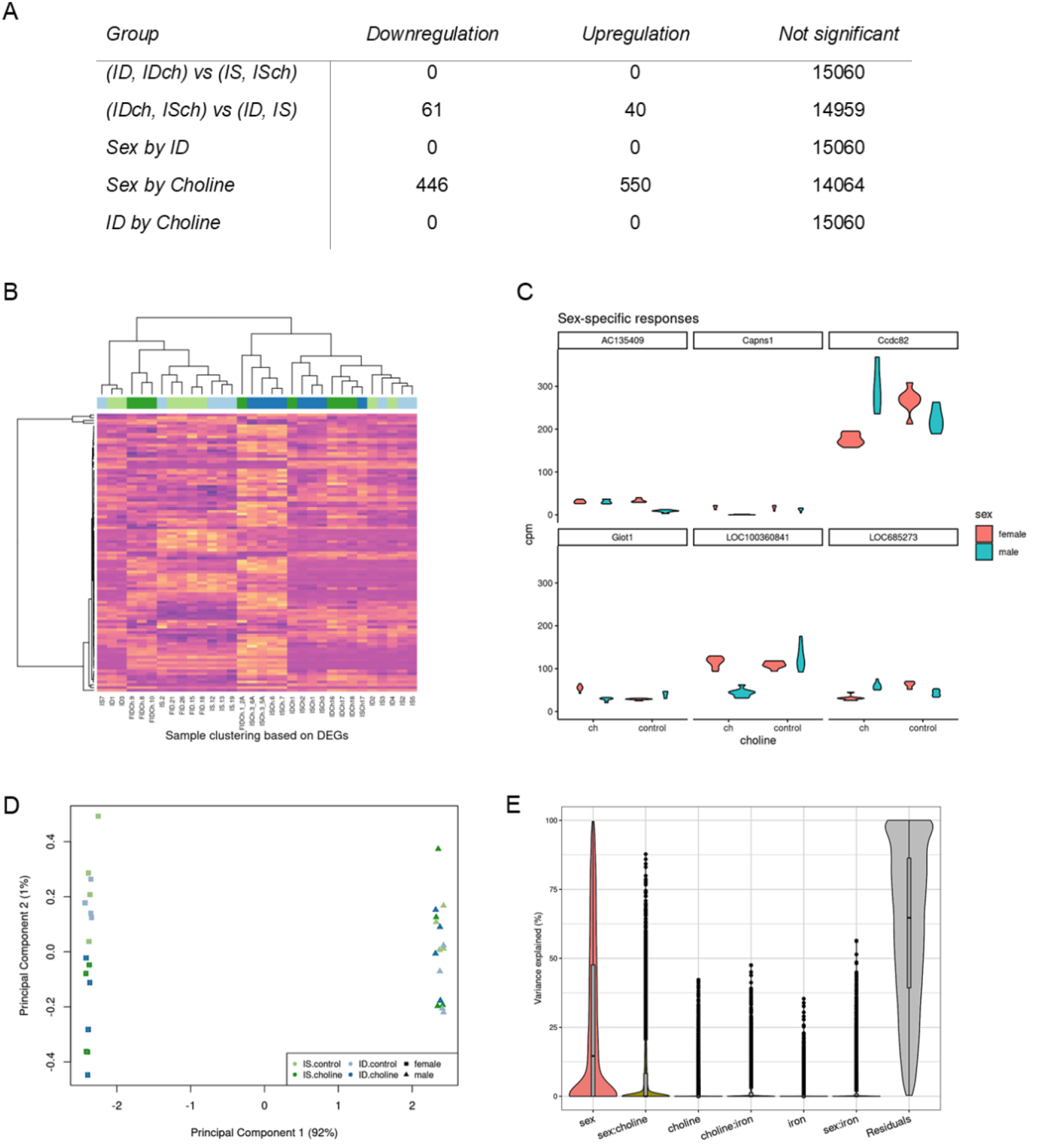
Bioinformatic integration of male and female RNA-seq datasets generated using different sequencing platforms. (A) Comparative analysis between groups showing a robust effect of sex-by-choline interaction on hippocampal gene expression. (B) Heatmap generated from unsupervised clustering showing a pattern that did not follow expected choline effects. (C) Top 6 representative genes showing sex-by-choline interaction effects. (D) Multidimensional plot showing a clear separation between sexes but not ID or choline treatment. (E) Analysis of variance revealed that variations in gene expression among combined libraries can be accounted by sex.

Effects of maternal choline supplementation: To estimate the effect of choline supplementation, the combined dataset was tested for DE genes between choline-supplemented (ISch, IDch, both sexes) vs choline non-supplemented (IS, ID, both sexes) groups. 101 genes (61 down- and 40 upregulated in choline-supplemented groups) showed significant changes (Figure 1A). Using these DE genes for an unsupervised clustering analysis of all samples, a resulting dendrogram did not show the expected trend of clustering following choline treatment (Figure 1B).

Sex-specific effects of choline supplementation: Analysis of sex and choline interaction showed a significant effect with 996 DE genes (Figure 1A). A violin plot of the top 6 genes with the lowest FDR value in the sex-choline interaction test showed differential effects of choline supplementation on gene profiles depending on the sex of the rats (Figure 1C). A similar plot of 6 randomly selected genes showed no such interaction (Supplemental Methods).

Assessing and managing confounding effects: Since male and female data were collected in different batches, we estimated the sex/batch contribution to variance and overall gene expression. Principle component analysis showed that sex/batch/technology can account for most of the variability in the combined dataset (Figure 1D). Analysis of variance showed that the variation across sex/batch/technology could account for a median of 14.6% of the variation in gene expression, while the median effects of choline, iron, and their interactions were negligible after correcting for all the others (Figure 1E). To remove the potential confounding effects, we followed up with separate analyses of each sex.

### Effects of maternal ID and choline supplementation in female offspring

Data from 16 female rats (4 from each treatment group) were analyzed. 14,965 of the 32,883 annotated genes passed the filtering process. Multidimensional reduction analysis showed a greater effect of choline supplementation than ID evidenced by a better group separation associated with choline than iron status (Figure 2A). Analysis of variance indicated that choline supplementation can explain about 20% of the variability in the expression profiles of all female animals (Figure 2B). Testing for DE genes showed little effects of iron status (Figure 2C, (ID, IDch) vs (IS, ISch)), indicating no considerable difference between groups that can be explained by iron status. Conversely, choline supplementation showed a clear effect on the expression changes of 1340 genes (629 down- and 711 upregulated) regardless of their iron status (Figure 2C, (IDch, ISch) vs (ID, IS)). Further assessment for the effect of choline supplementation on the IS control group showed 153 down- and 243 upregulated genes (Figure 2C, ISch vs IS). The clustering heatmap showed that the expression of most DE genes follows the choline treatment (Figure 2D). Analysis of functional changes using IPA to annotate these DE genes indicates reduced anxiety and other emotional behaviors, paired-pulse facilitation of synapse, and potentially viability of neuroglia (Z-score = −1.13), accompanied by increased migration of cerebral cortical cells and motor function (Figure 2E).

**Figure 2:**
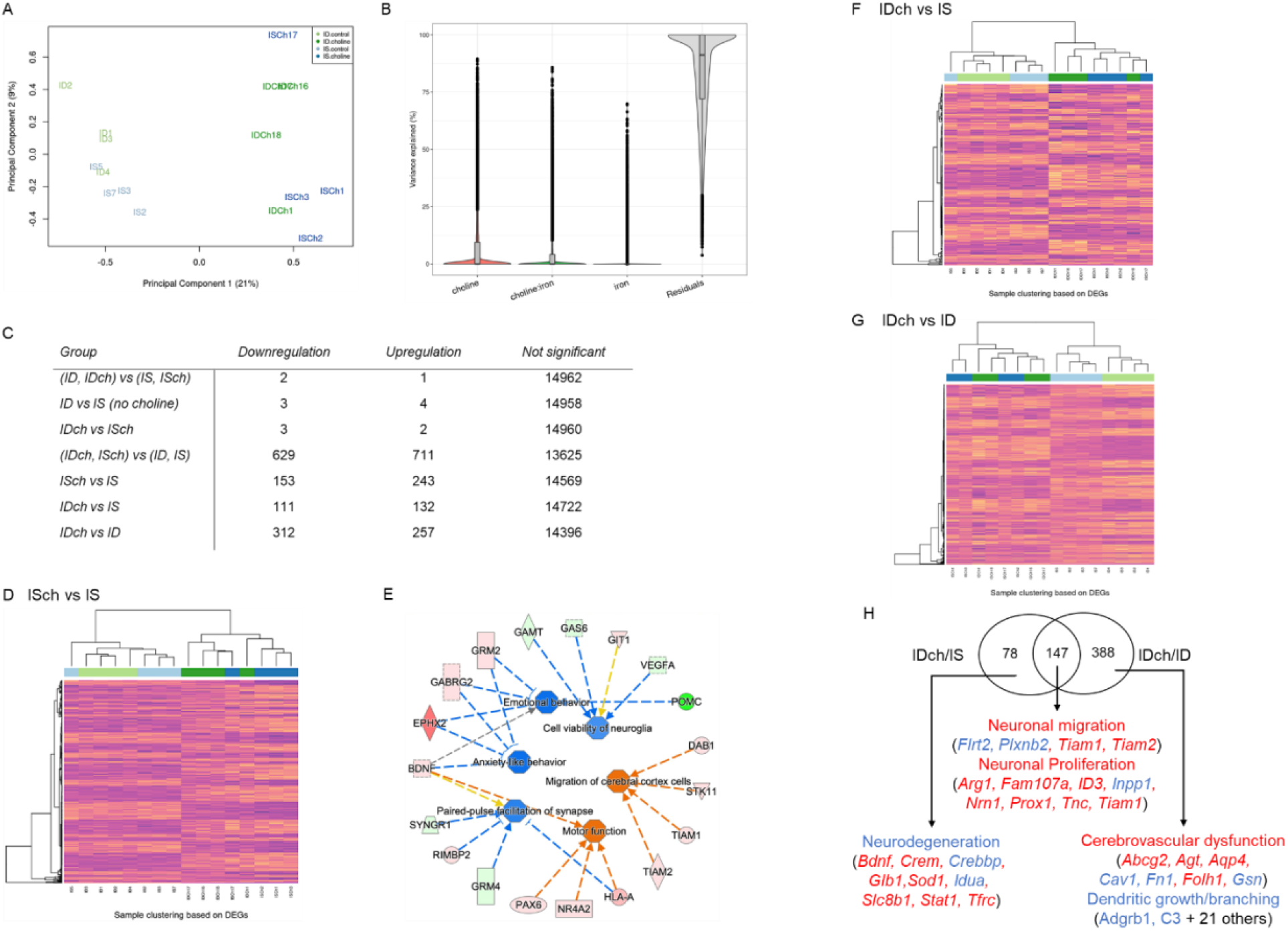
Effects of choline supplementation and ID on female hippocampal transcriptome. (A) Multidimensional plot showing a clear separation between choline-supplemented and non-supplemented groups, but not with iron status. (B) Analysis of variance showing that choline and choline-by-iron interaction can account for changes in gene expression. (C) Table showing gene expression changes among comparison groups. (D) Heatmap of DE genes from ISch vs IS comparison showing clear clustering between choline-supplemented and non-supplemented groups. (E) IPA annotated functional changes based on DE genes between ISch vs IS comparison. Blue = Inhibition, Orange = Activation, Yellow = Finding inconsistent with state of downstream molecule, Grey = Finding not predicted, Solid and Dashed arrows = direct and indirect interactions, Green-to-Red = Downregulated-to-Upregulated genes. (F-G) Heatmaps showing distinct clustering of choline treated samples using DE genes in IDch vs IS or ID groups. (H) IPA analysis of overlapping DE genes between IDch vs IS and IDch vs ID groups showing the rescue effects of choline treatment on specific biological activities.

To assess the rescue effects of choline supplementation to fetal-neonatal ID rats, IDch females were compared to IS females. 243 (111 down- and 132 upregulated) DE genes were identified between the two groups (Figure 2C, IDch vs IS). A hierarchical clustering heatmap showed a clear separation between choline supplemented compared to non-supplemented groups (Figure 2F). To further identify the interaction between ID and choline supplementation, libraries were compared between IDch and ID groups. 569 (312 down- and 257 upregulated) DE genes were found between groups (Figure 1C, IDch vs ID). Clustering using only DE genes with all libraries showed that samples were separated by choline supplementation status (Figure 2G). Within the choline non-supplemented groups, samples were also separated depending on their iron status; in contrast, the choline group samples were not clustered following the iron status (Figure 2G). The findings indicate that choline supplementation rescued ID by normalizing gene expression in the IDch group to that in the ISch group. Comparative analysis using IPA found 147 overlap-DE genes between IDch vs IS and IDch vs ID groups, which indicated activation of neuronal proliferation and migration (Figure 2H). Analysis of the non-overlap-DE genes showed that IDch group exhibits inhibition of neurodegeneration compared to the IS group, whereas activation of cerebrovascular dysfunction and inhibition of dendritic growth compared to the ID group (Figure 2H).

### Effects of maternal ID and choline supplementation in male offspring

Data from 16 male rats (4 from each treatment group) were analyzed. 14,305 of the 32,883 annotated genes passed the filtering process. Clustering analysis by multidimensional scaling plot showed that the principal sources of variation among samples do not correspond with iron or choline status. Choline supplementation had a stronger effect on the expression profiles than ID evidenced by a greater heterogeneity in the choline supplemented groups while the IS and ID samples showed less separation (Figure 3A). Testing for DE genes showed no effect of iron status (Figure 3B, (ID, IDch) vs (IS, ISch)). Comparison between choline supplemented vs non-supplemented groups identified 132 DE genes (Figure 3B, (IDch, ISch) vs (ID, IS)), which did not show the expected clustering based on dietary treatments (Figure 3C). Further comparative analysis of ISch vs IS groups showed 37 DE genes that were clustered based on choline status, indicating an effect of choline (Figure 3D). The rescue effects of choline supplementation were analyzed by comparing IDch to IS or ID groups. These analyses found 54 (IDch vs IS) and 276 (IDch vs ID) DE genes (Figure 3B). Clustering analyses of DE genes in the IDch vs IS comparison showed a clear separation between choline supplemented vs non-supplemented groups (Figure 3E), but not in the IDch vs ID comparison (data not shown). These findings indicate a larger effect of choline supplementation than iron status on long-term gene expression.

**Figure 3:**
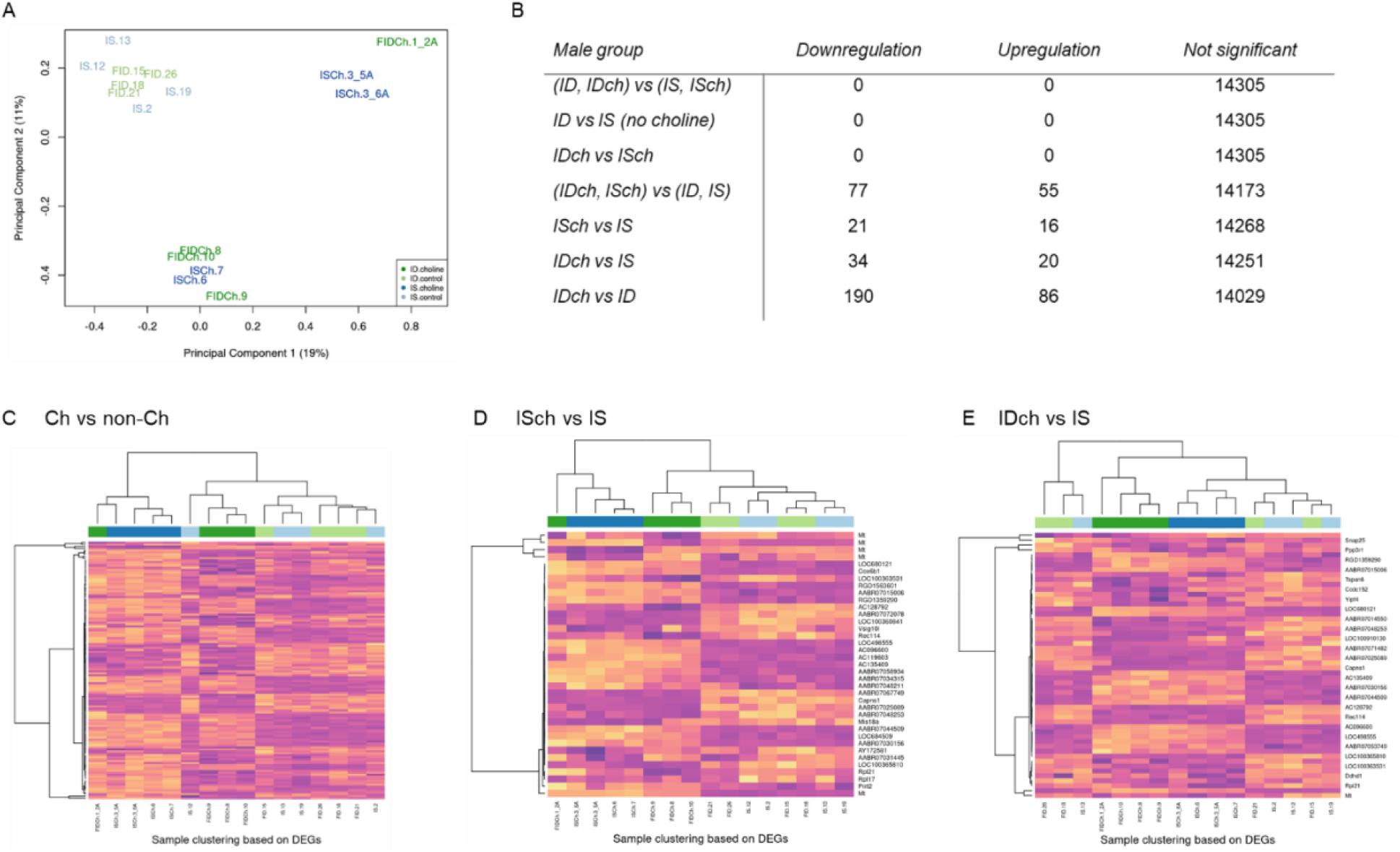
Effects of choline supplementation and ID on male hippocampal transcriptome. (A) Multidimensional plot showing that the sources of variation among samples do not correspond with either iron status or choline treatment. Choline supplementation seems to have a stronger effect than iron status on gene expression profiles as choline non-treated samples clustered together. (B) Table showing gene expression changes among comparison groups with the most effect in the IDch vs ID comparison. (C-E) Heatmaps from DE genes among comparison groups showing clear clustering among choline treated IS samples but less clear in the ID samples.

### Alternative approach

Given the genome-wide variance among the samples from the combined RNA-seq dataset producing too few DE genes defined by the FDR (<0.05) threshold, we used two additional approaches, Overrepresentation Analysis and Gene Set Enrichment Analysis, to glean at the altered biological functions attributable to sex, choline, iron, and their interactions.

### Overrepresentation Analysis

The top 300 genes identified by VP analysis were tested for overrepresentation of GO terms. The joint dataset combining male and female rats showed a statistically significant overrepresentation of GO terms associated with synaptic membrane and membrane transporter complex in the group of genes with highest variance explained by the choline-iron interaction (Figure 4A). These are consistent with the male data (Figure 4B); however, female rats showed a lower number of enriched GO terms (Figure 4C). Altogether, these findings suggested an interaction between choline supplementation and iron status in adult rat hippocampus, which is more pronounced in males than females. Volcano plots showed that most, if not all, of these genes have a significant p-value (<0.05, Figure 4D-F). IPA analysis of the top 300 VP outlier genes from the joint dataset indicated that choline significantly (absolute z ≥ 2.0) activated the transcriptional regulator cAMP responsive element binding protein (CREB1) gene network, which was inhibited among the genes regulated by ID-sex interaction (Figure 4G). The glutamate ionotropic receptor NMDA type subunit 3A (GRIN3A) gene network was inhibited among the genes regulated by ID-Ch interaction. Among genes affected by choline-sex interaction, inflammatory regulator transforming growth factor beta 1 (TGFb1) gene network was inhibited (Figure 4G). In males, ID activated the transcription factor fifth Ewing variant (FEV) gene network, which was mitigated by choline treatment (Figure 4H). In females, IPA analysis showed altered gene regulation indicating a propensity for inhibition of alpha synuclein (SNCA) gene network (z = −1.63) by choline supplementation (Ch), an inhibition of an intracellular cholesterol transporter Niemann-Pick C1 protein (NPC1) by ID-choline interaction, and an activation of the G protein-couple receptor adenosine A2a receptor (ADORA2A) gene network (z = 1.98) by ID (Figure 4I).

**Figure 4:**
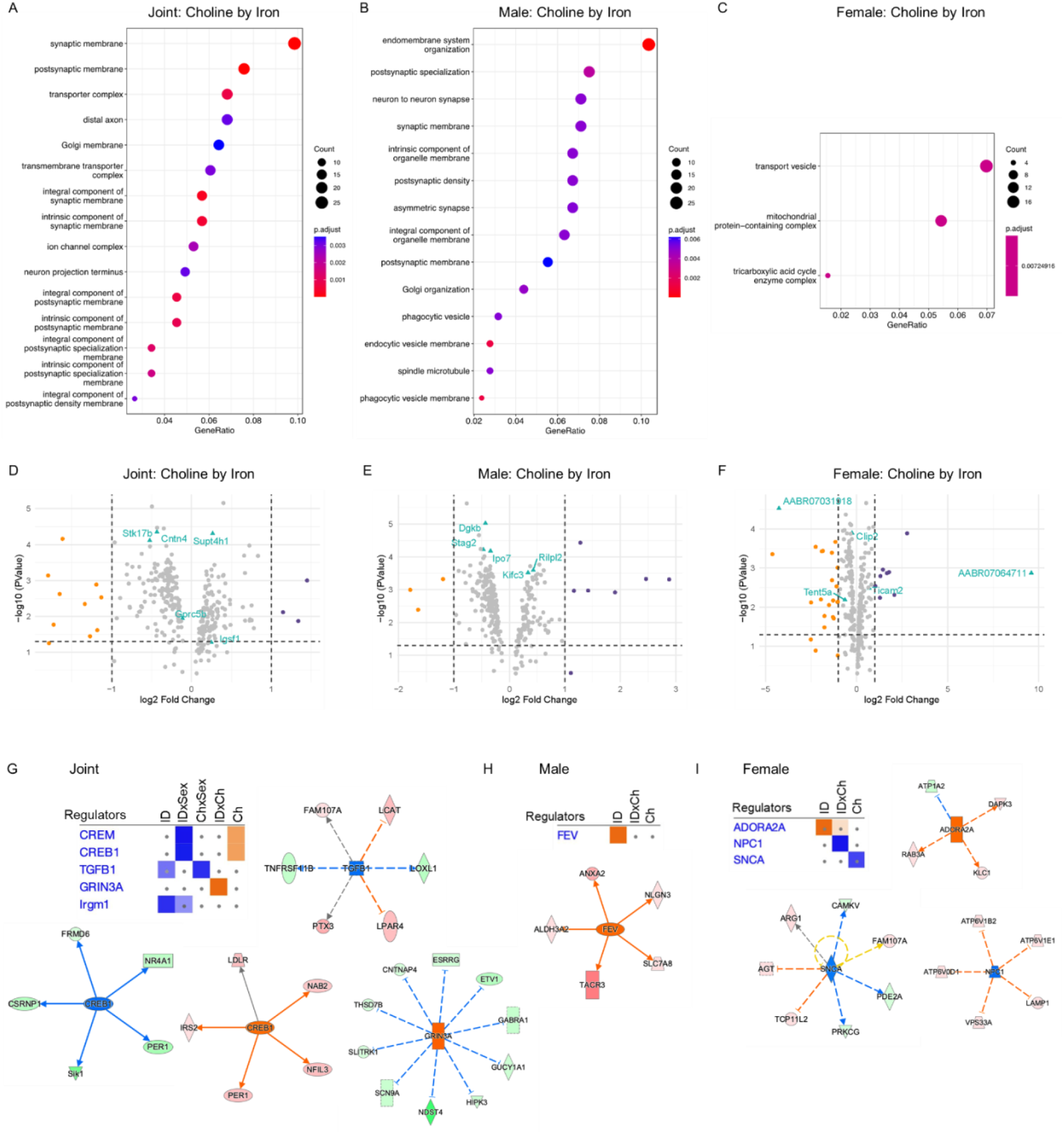
Alternative approach using top 300 VP outlier genes. (A-C) ORA analysis showing the effects of choline-by-iron interaction using sex-combined (A), male (B), and female (C) datasets. (D-F) Volcano plots showing genes with significant p-value (<0.05). (G-I) IPA-predicted changes of upstream regulators in the hippocampus using sex-combined (G), male (H), and female (I) datasets.

### Gene Set Enrichment Analysis

#### Effects of maternal choline and ID

Choline-driven gene expression profiles showed more suppressed than activated gene networks (Figure 5A, Choline). Sex was also a co-factor for gene expression variation as evidenced by a reversal of specific biological activities (Figure 5A, Sex by choline). Specifically, gene functions associated with epithelial-mesenchymal transition, apical junction and apical surface were downregulated in choline-supplemented females compared to males. On the other hand, gene functions associated with xenobiotic metabolism, fatty acid metabolism, and MYC targets V1 were upregulated in choline-supplemented females compared to males. ID-driven gene expression changes indicated more suppressed functions that were also influenced by sex. While the common trend for both sexes is a downregulation of pathways when fed with an ID diet (e.g., oxidative phosphorylation, DNA repair, ROS pathway; Figure 5A, ID), the sex-by-ID interaction test presents evidence of sex-specific effects (Figure 5A, Sex by ID). Particularly, MYC targets V1, oxidative phosphorylation, fatty acid metabolism, protein secretion and interferon alpha response are upregulated in ID females when compared to ID males, indicating an interaction of sex and iron status. Volcano plots showed genes with a significant p-value (<0.05, Figure 5B).

**Figure 5:**
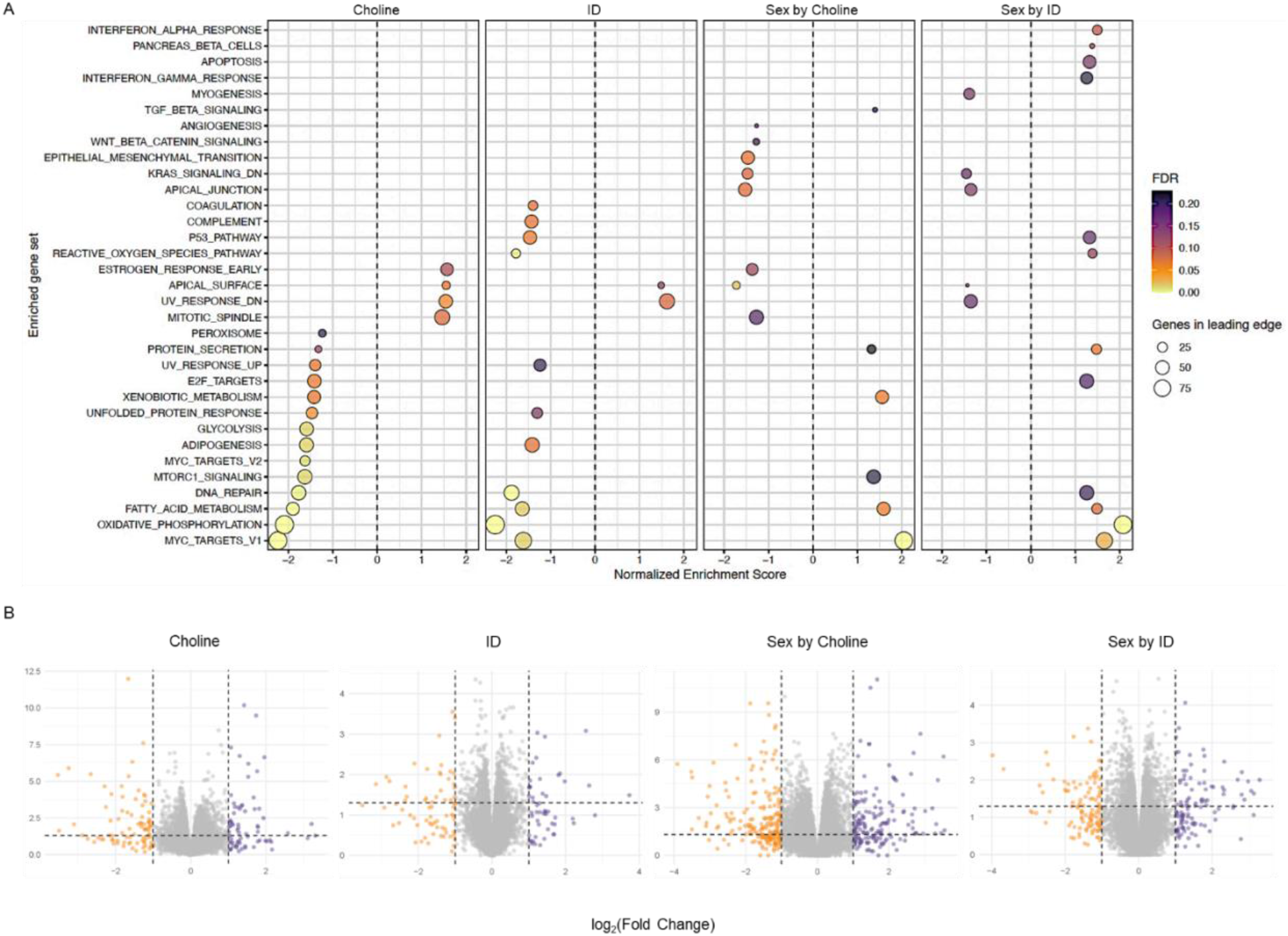
Alternative approach using ranked gene lists. (A) GSEA analysis using sex-combined datasets showing the effects of choline supplementation, ID, and their interaction with sex on enriched gene sets. Test were made using males, IS and non-choline supplemented diets as baseline. (B) Volcano plots showing genes with significant p-value (<0.05).

#### Sex-specific effects of prenatal choline and fetal-neonatal ID

To confirm the sex-specific effects, choline- and ID-driven DE genes were analyzed separately by sex. In females, GSEA analysis showed that choline supplementation induced changes in gene expression that indicate more activated biological functions, whereas ID resulted in more suppressed functions (Figure 6A). Notable changes include activation of interferon gamma (INFg) and tumor necrosis factor alpha (TNFa) signaling in choline-supplemented groups, and activation of INFa but inhibition of TNFa in the ID groups (Figure 6A). In contrast to the effects on females, choline- and ID-driven gene expression changes showed fewer significantly altered biological functions in males, where there were more suppressed activities (e.g., protein secretion, fatty acid metabolism) in the choline groups (ISch, IDch) but more activated activities (e.g., INFg, epithelial-mesenchymal transition) in the ID group (Figure 6B). In sum, results from the sex-separate analysis correlated with those from the joint GSEA, confirming the differential effects of choline supplementation and iron deficiency in specific pathways for female and male rats.

**Figure 6:**
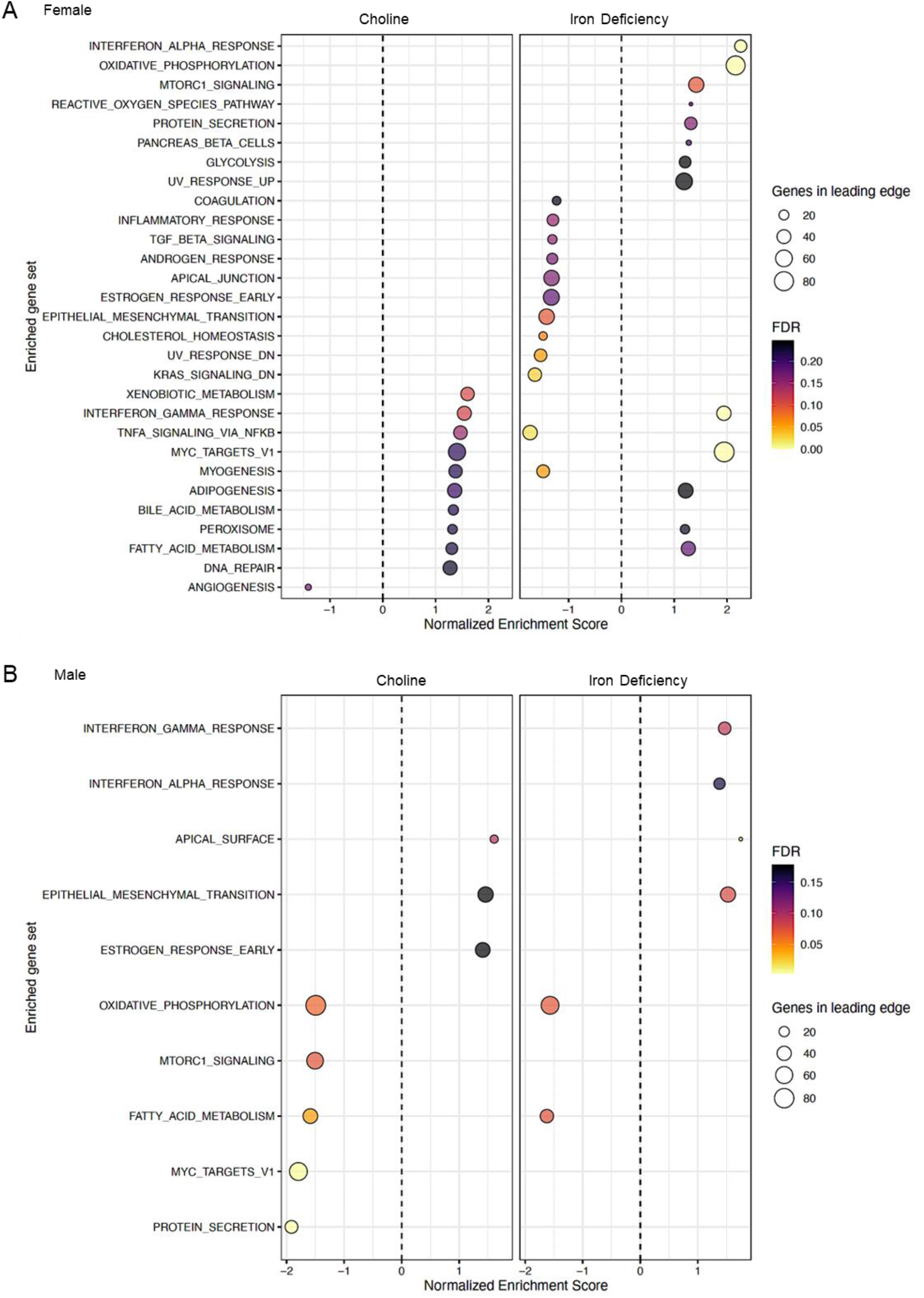
Alternative approach using ranked gene lists. GSEA plots showing the effects of choline supplementation and ID on enriched gene sets of female (A) and male (B) rats.

### Biomarkers

Using biomarker library available in the IPA database, ID- and choline-driven candidate genes identified by VP analysis (top 300 genes) were assessed for potential roles as markers of long-term effects (Table 1). In female rats, both ID, choline supplementation, and their interaction changed expression of markers regulating inflammation (*Arg1, Cxcr4, Cx3cr1*, and *Pla2g7*). In male rats, altered expression of notable markers include regulators of cellular growth and survival (*Anxa2, Eno2, Nrg1*) induced by ID, protein secretion (*Inhbb*) induced by choline supplementation (Ch); and synaptogenesis (*Pten, Pias1*) induced by ID-by-choline interaction.

**Table 1:**
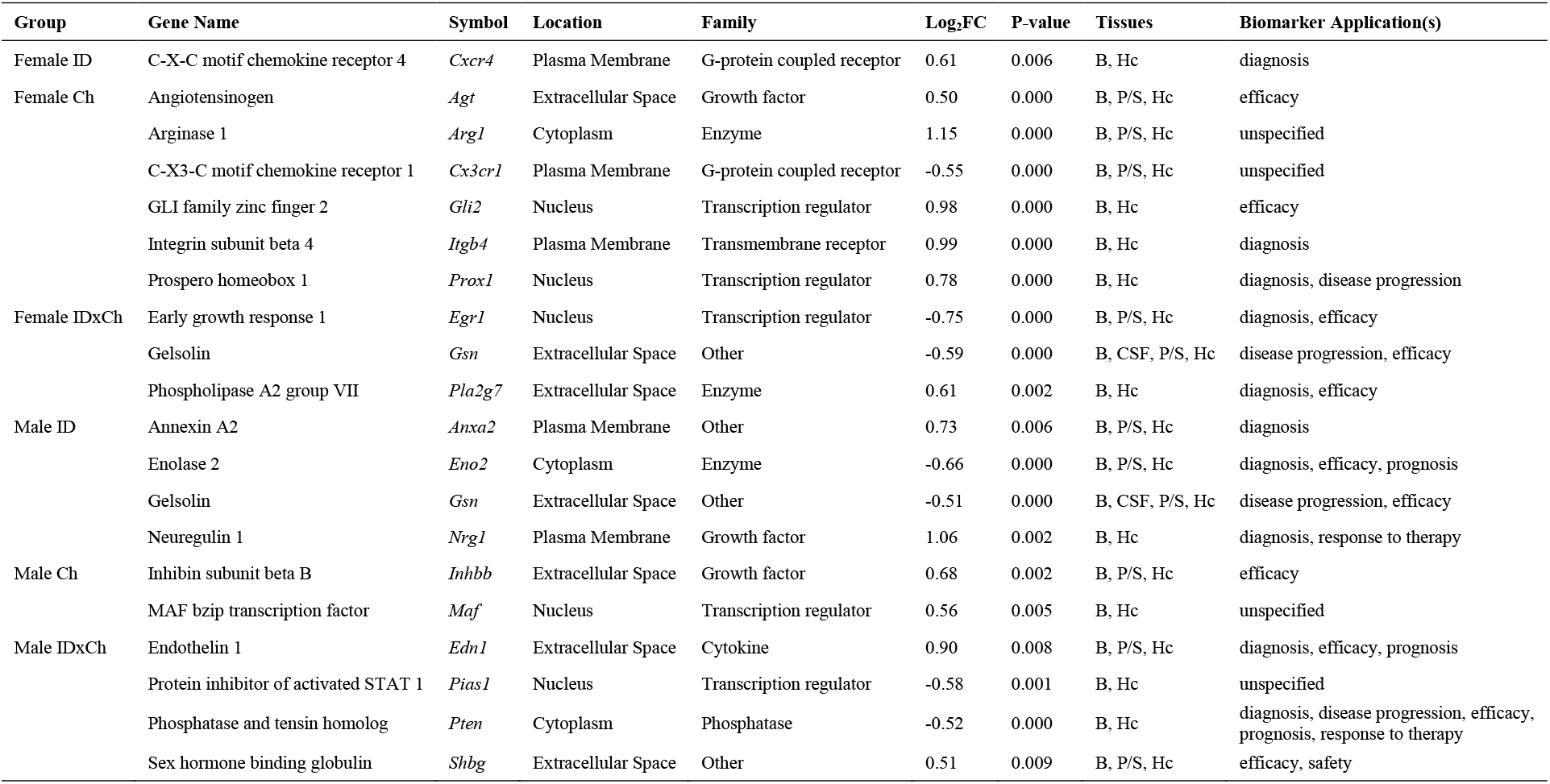
IPA-mapped known biomarkers found in humans, mice and rats. Selection criteria include absolute log_2_FC ≥ 0.50, p <0.05. Abbreviations: Blood (B), Cerebrospinal fluid (CSF), Plasma/Serum (P/S), and Hippocampus (Hc)

## Discussions and Conclusions

By assessing global changes in hippocampal gene expression, this study presents novel findings pertaining to long-term sex-specific hippocampal effects of early-life ID and prenatal choline supplementation on gene regulation and changes in pre-defined neurologic biomarkers. DE genes identified by the traditional approach with FDR value <0.05 showed that major factors contributing to the long-term changes in hippocampal gene regulation were sex, choline supplementation, and iron status in that order. Interaction between sex and choline supplementation showed the most robust effects compared to the iron status by sex interaction or individual factors on hippocampal gene expression. Overall, female rats exhibited more effects of choline supplementation and ID than male rats. Because of the high variations among the combined samples due to integration from two separate RNA-seq batches using two different sequencing technologies, alternative approaches were applied to identify genes with variance driven by the analyzed variables (ID, choline, and their interaction) using genes deviated from the genome-wide trend [27], as well as to use ranked gene lists to identify altered gene networks by GSEA analysis. Both ID and choline supplementation showed similar patterns of sex-specific effects on changes in hippocampal gene networks and associated biological processes.

Little is known about long-term effects of prenatal choline supplementation on the rat hippocampal transcriptome. Here we showed that prenatal choline supplementation induced long-term sex-specific gene expression changes in adult hippocampal transcriptomes with greater effects in female than male rats. The robust effects in female rats suggest a more amenable epigenetic regulation, which indicates a more adaptable developing hippocampus in female than male rats. This adaptability may confer sex-specific advantages such as neurodevelopmental resilience in advent of prenatal adverse exposures (e.g., nutritional deficiencies, maternal stress) [28, 29]. In addition, prenatal choline supplementation produced more significant transcriptomic changes in adult rat hippocampus compared to fetal-neonatal ID in either sex, implicating an important role of epigenetic regulation in the backdrop of ID, given that choline is a known methyl donor that influences epigenetic modifications [30]. These findings could also provide a molecular explanation for the recovery of ID-induced impairments of neuronal structure and cognition even during the period of limited iron substrate [25, 26, 31].

Our study further expands the knowledge of the interaction between choline and iron status previously discovered [21, 26, 32] with demonstration of sex-specific effects. In ISch females, choline supplementation might provide beneficial effects by altering expression of genes that regulate emotions including anxiety and motor function. Choline could also reprogram anti-inflammatory responses with an activation of INFg and TNFa [33, 34] in females (Figure 6). In this regard, there is evidence that prenatal choline supplementation mitigates fetal alcohol-exposure induced pro-inflammatory responses in the adult rat hippocampus [35]. Thus, the role of prenatal choline supplementation in neuroinflammation deserves further analyses. On the other hand, choline supplementation could also produce undesired effects. In the ISch males, choline supplementation might induce adverse cellular effects similar to those observed in iron-deficient animals by suppressing the activity of oxidative phosphorylation and fatty acid metabolism [36–38] as well as activating cell adhesion and migration by increasing epithelial-mesenchymal transition. These findings indicate both benefits and potential risks of prenatal choline supplementation to the IS rats. Thus, the use of choline as a therapeutic agent needs additional investigation, particularly in the long-term gene expression changes that modulate neurobehavioral functions [26].

Regarding the rescue effects of prenatal choline supplementation on ID-altered gene expression, a greater number of genes was found when compared to ID than IS hippocampus (IDch vs ID compared to IDch vs IS) in both sexes, suggesting a robust rescue effect of choline supplementation on gene regulation, particularly among genes regulating MTOR and CREB signaling and inflammatory response. These effects of ID have been previously reported in models of perinatal ID [8, 39, 40]. Novel rescue effects of choline include normalization of ID-activated ADORA2 gene network in females and FEV gene network in males. ADORA2 gene network is implicated in emotional behaviors and neurodegenerative disorders [41, 42]. FEV gene network is implicated in depression and anxiety [43]. It is worthwhile to note the suppressing effect of choline on alpha-synuclein (SNCA), whose translation is regulated by the iron-response element (IRE) in the 5’-UTR of its transcript and its overexpression is associated with neuropathologies such as Parkinson’s and Alzheimer’s diseases [44–47]. The finding suggests a potential mechanism for the neuroprotective effect of prenatal choline supplementation. Not all ID-induced effects (e.g., activated adipogenesis in females and suppressed oxidative phosphorylation in males) could be rescued by choline supplementation, suggesting the limitation of choline treatment.

Finally, a potentially important finding of the present study is the identification of biomarkers that are readily accessible in bodily fluids. Long-term specific changes due to ID (e.g., *Nrg1*), sex (e.g., *Cxcr4, Eno2*), or choline treatment (e.g., *Arg1, Pias1*) in the rat hippocampus indicate that these genes can be utilized as potential markers to gauge the health status of the hippocampus as well as other brain regions. Future studies can determine the relationship of these markers between blood and brain compartments following ID anemia. In this regard, our PIA model [8] would be an ideal system to determine the utility of these biomarkers as we could generate various degrees of ID anemia severity (i.e., 25 vs 18% hematocrit). Determining the predictive values of these potential biomarkers from an easily accessible source (e.g., blood) would be critically important in the prevention and treatment of brain ID in order to reduce the life-long risks of poor neurobehavioral outcomes.

In conclusion, the power of the present study is in the global unbiased approach to identify differential gene expression changes due to ID, sex, and choline treatment. The long-term changes in gene expression, and their regulation, encompass various biological processes (e.g., synaptogenesis, inflammation). The present study has achieved our immediate goals in uncovering sex-specific effects of early-life ID and prenatal choline treatment on gene expression, laying down the groundwork by providing targets for future studies to address the causal relationship between these specific gene expression changes and hippocampal-dependent behavioral outcomes as well as more broadly functional consequences of the cortical-hippocampal-striatal circuitry.

Sources of funding: This work was funded by NIH R01NS099178 to P.V.T. and R01HD29421 to M.K.G.

## Supporting information

Supplemental Method-Bioinformatics

## References

1. Georgieff MK: Iron deficiency in pregnancy. Am J Obstet Gynecol 2020, 223(4):516–524.

2. Insel BJ, Schaefer CA, McKeague IW, Susser ES, Brown AS: Maternal iron deficiency and the risk of schizophrenia in offspring. Arch Gen Psychiatry 2008, 65(10):1136–1144.

3. Wiegersma AM, Dalman C, Lee BK, Karlsson H, Gardner RM: Association of Prenatal Maternal Anemia With Neurodevelopmental Disorders. JAMA Psychiatry 2019, 76(12):1294–1304.

4. Youdim MB: Brain iron deficiency and excess; cognitive impairment and neurodegeneration with involvement of striatum and hippocampus. Neurotox Res 2008, 14(1):45–56.

5. McClorry S, Zavaleta N, Llanos A, Casapía M, Lönnerdal B, Slupsky CM: Anemia in infancy is associated with alterations in systemic metabolism and microbial structure and function in a sex-specific manner: an observational study. Am J Clin Nutr 2018, 108(6):1238–1248.

6. Nopoulos PC, Conrad AL, Bell EF, Strauss RG, Widness JA, Magnotta VA, Zimmerman MB, Georgieff MK, Lindgren SD, Richman LC: Long-term outcome of brain structure in premature infants: effects of liberal vs restricted red blood cell transfusions. Arch Pediatr Adolesc Med 2011, 165(5):443–450.

7. McCoy TE, Conrad AL, Richman LC, Brumbaugh JE, Magnotta VA, Bell EF, Nopoulos PC: The relationship between brain structure and cognition in transfused preterm children at school age. Dev Neuropsychol 2014, 39(3):226–232.

8. Singh G, Wallin DJ, Abrahante Lloréns JE, Tran PV, Feldman HA, Georgieff MK, Gisslen T: Dose-and sex-dependent effects of phlebotomy-induced anemia on the neonatal mouse hippocampal transcriptome. Pediatr Res 2021.

9. Rudy M, Mayer-Proschel M: Iron Deficiency Affects Seizure Susceptibility in a Time-and Sex-Specific Manner. ASN Neuro 2017, 9(6):1759091417746521.

10. Matveeva TM, Singh G, Gisslen TA, Gewirtz JC, Georgieff MK: Sex differences in adult social, cognitive, and affective behavioral deficits following neonatal phlebotomy-induced anemia in mice. Brain Behav 2021, 11(3):e01780.

11. Woodman AG, Noble RMN, Panahi S, Gragasin FS, Bourque SL: Perinatal iron deficiency combined with a high salt diet in adulthood causes sex-dependent vascular dysfunction in rats. J Physiol 2019, 597(18):4715–4728.

12. Cao C, Prado MA, Sun L, Rockowitz S, Sliz P, Paulo JA, Finley D, Fleming MD: Maternal Iron Deficiency Modulates Placental Transcriptome and Proteome in Mid-Gestation of Mouse Pregnancy. J Nutr 2021, 151(5):1073–1083.

13. Marell PS, Blohowiak SE, Evans MD, Georgieff MK, Kling PJ, Tran PV: Cord Blood-Derived Exosomal CNTN2 and BDNF: Potential Molecular Markers for Brain Health of Neonates at Risk for Iron Deficiency. Nutrients 2019, 11(10):2478.

14. Duarte-Guterman P, Yagi S, Chow C, Galea LA: Hippocampal learning, memory, and neurogenesis: Effects of sex and estrogens across the lifespan in adults. Horm Behav 2015, 74:37–52.

15. Zafer D, Aycan N, Ozaydin B, Kemanli P, Ferrazzano P, Levine JE, Cengiz P: Sex differences in Hippocampal Memory and Learning following Neonatal Brain Injury: Is There a Role for Estrogen Receptor-α? Neuroendocrinology 2019, 109(3):249–256.

16. Barks A, Hall AM, Tran PV, Georgieff MK: Iron as a model nutrient for understanding the nutritional origins of neuropsychiatric disease. Pediatr Res 2019, 85(2):176–182.

17. Callahan LS, Thibert KA, Wobken JD, Georgieff MK: Early-life iron deficiency anemia alters the development and long-term expression of parvalbumin and perineuronal nets in the rat hippocampus. Dev Neurosci 2013, 35(5):427–436.

18. Beard JL, Wiesinger JA, Connor JR: Pre- and postweaning iron deficiency alters myelination in Sprague-Dawley rats. Dev Neurosci 2003, 25(5):308–315.

19. DeMaman AS, Homem JM, Lachat JJ: Early iron deficiency produces persistent damage to visual tracts in Wistar rats. Nutr Neurosci 2008, 11(6):283–289.

20. Rao R, Tkac I, Townsend EL, Ennis K, Gruetter R, Georgieff MK: Perinatal iron deficiency predisposes the developing rat hippocampus to greater injury from mild to moderate hypoxia-ischemia. J Cereb Blood Flow Metab 2007, 27(4):729–740.

21. Liu SX, Barks AK, Lunos S, Gewirtz JC, Georgieff MK, Tran PV: Prenatal Iron Deficiency and Choline Supplementation Interact to Epigenetically Regulate Jarid1b and Bdnf in the Rat Hippocampus into Adulthood. Nutrients 2021, 13(12).

22. Krämer A, Green J, Pollard J, Jr., Tugendreich S: Causal analysis approaches in Ingenuity Pathway Analysis. Bioinformatics 2014, 30(4):523–530.

23. Ashburner M, Ball CA, Blake JA, Botstein D, Butler H, Cherry JM, Davis AP, Dolinski K, Dwight SS, Eppig JT et al: Gene ontology: tool for the unification of biology. The Gene Ontology Consortium. Nat Genet 2000, 25(1):25–29.

24. Liberzon A, Subramanian A, Pinchback R, Thorvaldsdottir H, Tamayo P, Mesirov JP: Molecular signatures database (MSigDB) 3.0. Bioinformatics 2011, 27(12):1739–1740.

25. Kennedy BC, Dimova JG, Siddappa AJ, Tran PV, Gewirtz JC, Georgieff MK: Prenatal choline supplementation ameliorates the long-term neurobehavioral effects of fetal-neonatal iron deficiency in rats. J Nutr 2014, 144(11):1858–1865.

26. Tran PV, Kennedy BC, Pisansky MT, Won KJ, Gewirtz JC, Simmons RA, Georgieff MK: Prenatal Choline Supplementation Diminishes Early-Life Iron Deficiency-Induced Reprogramming of Molecular Networks Associated with Behavioral Abnormalities in the Adult Rat Hippocampus. J Nutr 2016, 146(3):484–493.

27. Hoffman GE, Schadt EE: variancePartition: interpreting drivers of variation in complex gene expression studies. BMC Bioinformatics 2016, 17(1):483.

28. Russell JA, Brunton PJ: Giving a good start to a new life via maternal brain allostatic adaptations in pregnancy. Front Neuroendocrinol 2019, 53:100739.

29. Weinstock M: The potential influence of maternal stress hormones on development and mental health of the offspring. Brain Behav Immun 2005, 19(4):296–308.

30. Zeisel SH: Epigenetic mechanisms for nutrition determinants of later health outcomes. Am J Clin Nutr 2009, 89(5):1488S–1493S.

31. Bastian TW, von Hohenberg WC, Kaus OR, Lanier LM, Georgieff MK: Choline Supplementation Partially Restores Dendrite Structural Complexity in Developing Iron-Deficient Mouse Hippocampal Neurons. J Nutr 2022, 152(3):747–757.

32. Carter RC, Senekal M, Duggan CP, Dodge NC, Meintjes EM, Molteno CD, Jacobson JL, Jacobson SW: Gestational weight gain and dietary energy, iron, and choline intake predict severity of fetal alcohol growth restriction in a prospective birth cohort. Am J Clin Nutr 2022, 116(2):460–469.

33. Deckert-Schlüter M, Bluethmann H, Kaefer N, Rang A, Schlüter D: Interferon-gamma receptor-mediated but not tumor necrosis factor receptor type 1-or type 2-mediated signaling is crucial for the activation of cerebral blood vessel endothelial cells and microglia in murine Toxoplasma encephalitis. Am J Pathol 1999, 154(5):1549–1561.

34. Tau G, Rothman P: Biologic functions of the IFN-gamma receptors. Allergy 1999, 54(12):1233–1251.

35. Baker JA, Breit KR, Bodnar TS, Weinberg J, Thomas JD: Choline Supplementation Modifies the Effects of Developmental Alcohol Exposure on Immune Responses in Adult Rats. Nutrients 2022, 14(14).

36. Chung YJ, Swietach P, Curtis MK, Ball V, Robbins PA, Lakhal-Littleton S: Iron-Deficiency Anemia Results in Transcriptional and Metabolic Remodeling in the Heart Toward a Glycolytic Phenotype. Front Cardiovasc Med 2020, 7:616920.

37. Bastian TW, Rao R, Tran PV, Georgieff MK: The Effects of Early-Life Iron Deficiency on Brain Energy Metabolism. Neurosci Insights 2020, 15:2633105520935104.

38. Stangl GI, Kirchgessner M: Different degrees of moderate iron deficiency modulate lipid metabolism of rats. Lipids 1998, 33(9):889–895.

39. Fretham SJ, Carlson ES, Georgieff MK: Neuronal-specific iron deficiency dysregulates mammalian target of rapamycin signaling during hippocampal development in nonanemic genetic mouse models. J Nutr 2013, 143(3):260–266.

40. Barks A, Fretham SJB, Georgieff MK, Tran PV: Early-Life Neuronal-Specific Iron Deficiency Alters the Adult Mouse Hippocampal Transcriptome. J Nutr 2018, 148(10):1521–1528.

41. Oliveira S, Ardais AP, Bastos CR, Gazal M, Jansen K, de Mattos Souza L, da Silva RA, Kaster MP, Lara DR, Ghisleni G: Impact of genetic variations in ADORA2A gene on depression and symptoms: a cross-sectional population-based study. Purinergic Signal 2019, 15(1):37–44.

42. Siokas V, Mouliou DS, Liampas I, Aloizou AM, Folia V, Zoupa E, Papadimitriou A, Lavdas E, Bogdanos DP, Dardiotis E: Analysis of ADORA2A rs5760423 and CYP1A2 rs762551 Genetic Variants in Patients with Alzheimer’s Disease. Int J Mol Sci 2022, 23(22).

43. Albert PR, Vahid-Ansari F, Luckhart C: Serotonin-prefrontal cortical circuitry in anxiety and depression phenotypes: pivotal role of pre-and post-synaptic 5-HT1A receptor expression. Front Behav Neurosci 2014, 8:199.

44. Beyer K, Ariza A: alpha-Synuclein posttranslational modification and alternative splicing as a trigger for neurodegeneration. Mol Neurobiol 2013, 47(2):509–524.

45. Courte J, Bousset L, Boxberg YV, Villard C, Melki R, Peyrin JM: The expression level of alpha-synuclein in different neuronal populations is the primary determinant of its prion-like seeding. Sci Rep 2020, 10(1):4895.

46. Febbraro F, Giorgi M, Caldarola S, Loreni F, Romero-Ramos M: alpha-Synuclein expression is modulated at the translational level by iron. Neuroreport 2012, 23(9):576–580.

47. Zhou ZD, Tan EK: Iron regulatory protein (IRP)-iron responsive element (IRE) signaling pathway in human neurodegenerative diseases. Mol Neurodegener 2017, 12(1):75.

