## Supplemental Method-Bioinformatics for "Sex-Specific Effects of Early-Life Iron Deficiency and Prenatal Choline Treatment on Adult Rat Hippocampal Transcriptome"

Experimental approach
Analytical hypothesis
Expected sources of variation
Analysis details
Joint Analysis
Analysis of Female Pups
Analysis of Male Pups
Conclusions
GSEA and ORA notes

### Effect of Iron Deficiency and Choline Supplementation in the Hippocampal Transcriptomes of Rat Pups

Natalia Calixto Mancipe

2022-12-21

#### Experimental approach

A total of 32 rats were included in this experiment. The sampled rats are pups from maternal individuals subject to one of four diet regimens: iron sufficient diet (IS.control), iron deficient diet (ID.control), iron sufficient with choline supplementation (IS.choline), or iron deficient with choline supplementation (ID.choline). Litters were culled to eight pups of equal sex numbers at birth (four female pups and four male pups). RNA was isolated from the hippocampal lobes of the pups.

RNA from female pups and male pups were extracted and sequenced in separate batches. Female pups were sequenced with 150bp paired end reads on a NovaSeq. Male pups were sequenced with 50bp paired end reads on a HiSeq2000. Data from the male pups were previously analyzed in a 2016 publication (PMID: 26865644), and data from the female pups are being analyzed for a manuscript in preparation (preprint: https://www.biorxiv.org/content/10.1101/2022.06.29.498159v1 (https://www.biorxiv.org/content/10.1101/2022.06.29.498159v1)).

#### Analytical hypothesis

Genes underlying the response to maternal iron deficiency and choline supplementation have both sex-shared and sex specific responses. The sex-specific responses will contribute to differential outcomes of iron deficiency and choline supplementation between the sexes. Because the data were collected in two batches that are perfectly confounded with sex and sequencing technology, the sex-diet interaction effects are also perfectly confounded with batch and technology, thus need further analysis.

#### Expected sources of variation

- Sex (perfectly confounded with batch and technology)
- Individual genetic variation
- Diet (iron status, choline supplementation and interaction between both factors)

#### Analysis details

- **Filtering and normalization**: Gene raw counts were filtered to include only genes at least 300 bp long and that are expressed in at least one of the four dietary groups.
- **Statistical model**: Expression values were modeled by a generalized linear model (glm) with a negative binomial distribution using either the dietary group as explanatory variable if the model was built for each sex separately (16 RNA-seq libraries), or the combination of iron, choline and sex and their interaction as explanatory variables, using all 32 RNA-Seq libraries. For the analysis of variance a linear mixed model with sex, choline and iron and their interactions was used.
- **Analysis**: Differential gene expression was tested using a quasi-likelihood F test on the fitted glms and the following contrasts.
  - Reviewer suggestions using a full model (32 samples):
    - ID rats vs IS rats
    - Choline-supplemented vs control
    - Interaction sex:iron
    - Interaction sex:choline
  - Defined by researcher:
    - Iron deficiency (ID.control vs IS.control in male and female rats separately)
    - Iron deficiency and choline supplementation (ID.choline vs IS.choline in male and female rats separately)
    - Choline supplementation (IS.choline vs IS.control in male and female rats separately)
    - Rescued effects of choline supplementation (ID.choline vs. IS.control in male and female rats separately)
  - Additional tests:
    - Effect of choline supplementation in ID diets (ID.choline vs ID.control for each sex separatedly)
- **Diferentially expressed genes**: Genes with an FDR < 0.05 were taken as differentially expressed between the groups compared. These genes were used to perform an unsupervised hierarchical clustering of the samples and a heatmap depicting the counts per million (cpm) values of the DEGs in each group. The full list of test results for each comparison can be found in the .deg.txt files.

#### Joint Analysis

Data from female and male rats were combined and used to model gene expression. After filtering short and lowly expressed genes, the analysis set contained 15390 loci.

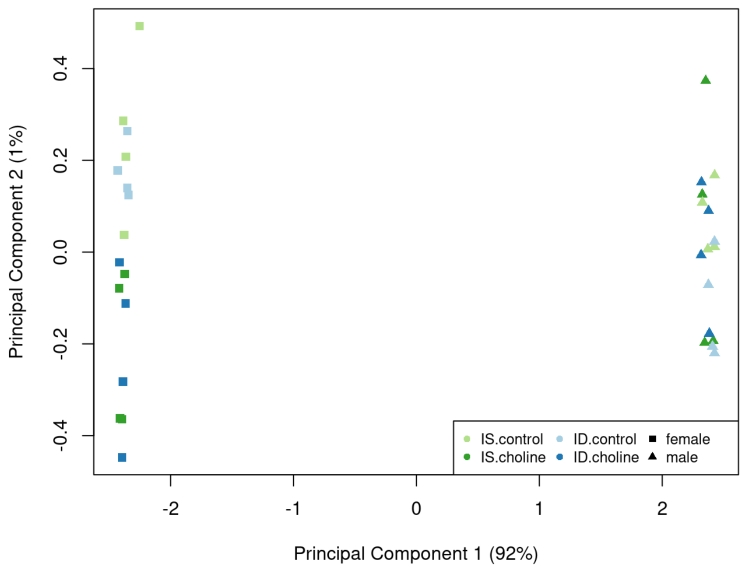


```
## png
##   2
```

The exploratory MDS plot shows that the PC1 correlates with the sex/batch group and can explain most of the variability in the data.

#### Analysis of variance

```
## Saving 7 x 6 in image
```

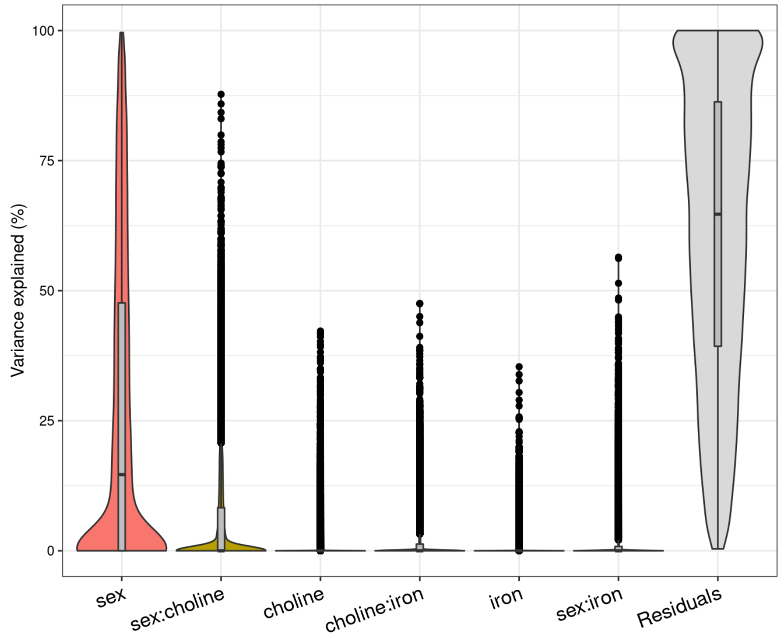


When both datasets are examined together with a linear mixed model the gene expression can not be successfully modeled using the explanatory variables of the study (sex, iron, choline and their interactions) as shown in the genome wide median variance plot; i.e. most of the genes do not to respond to these variables. The variation across sex/batch/technology explains a median of 14.6 % of the variation in gene expression, while the median effects of choline, iron and their interactions are negligible after correcting for all the others.

The top 100 genes that deviate from the genome-wide trend (outliers from the variance plot above) are selected and stored in the vp.txt files, the variance partition of the top 10 genes for each variable are plotted and saved as vp.pdf files.

```
## Saving 7 x 6 in image
## Saving 7 x 6 in image
## Saving 7 x 6 in image
## Saving 7 x 6 in image
## Saving 7 x 6 in image
## Saving 7 x 6 in image
```

#### Differential gene expression

To estimate the effect of dietary iron levels, choline supplementation and sex, the expression profiles of each gene were modeled with a generalized linear model using the edgeR package as y = f(sex + iron + choline + iron:sex + choline:sex + iron:choline). The threshold for differential expression is defined as FDR < 0.05.

##### Effect of iron deficiency

```
##           ID
## Down       0
## NotSig 15060
## Up         0
```

No genes were found differentially expressed as a response to ID.

##### Effect of choline supplementation

```
##        choline
## Down        61
## NotSig   14959
## Up          40
```

| | SYMBOL | FDR | logFC | DESCRIPTION |
|---|---|---|---|---|
| 15745 | LOC100360841 | 0.0000000 | -1.6636339 | ribosomal protein L37 |
| 19711 | AABR07044509 | 0.0000005 | 1.4192640 | NULL |
| 19561 | Capns1 | 0.0000012 | -4.5820674 | calpain small subunit 1-like |
| 16696 | AC135409 | 0.0000012 | 1.7397002 | NULL |
| 5197 | LOC680121 | 0.0000101 | 0.7404338 | similar to heat shock protein 8 |
| 18038 | LOC100365810 | 0.0000630 | -1.2634996 | 40S ribosomal protein S17-like |
| 16516 | LOC498555 | 0.0000742 | 0.8089468 | similar to 60S acidic ribosomal protein P2 |
| 21050 | LOC684509 | 0.0000937 | 1.0688084 | similar to NADH-ubiquinone oxidoreductase B9 subunit (Complex I-B9) (CI-B9) |
| 4417 | AABR07049516 | 0.0001713 | 0.8488493 | NULL |
| 17683 | LOC103692785 | 0.0001713 | -0.4335435 | 60S ribosomal protein L39 |

Choline supplemented diets did show a statistically significant effect in the expression profiles of 124 genes. Using these DEGs to cluster all samples results in a dendogram that does not follow the expected trend of clustering due to choline status.

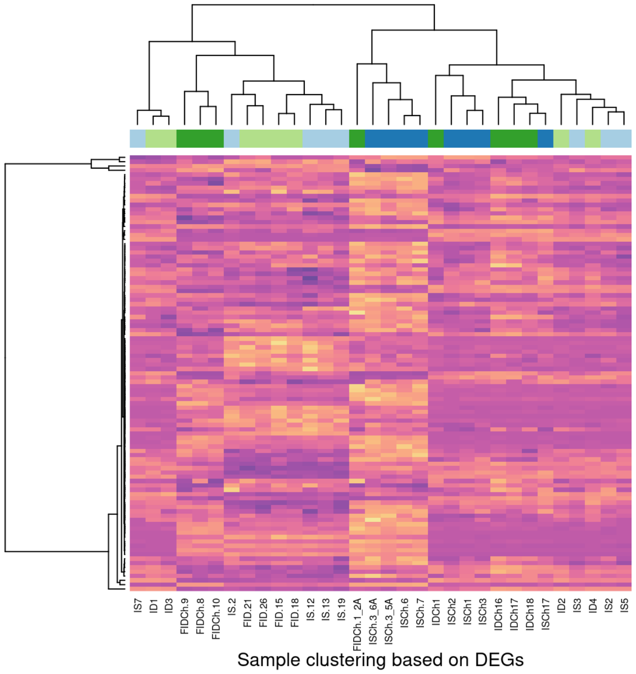


```
## png
##   2
```

##### Sex/batch specific responses to iron deficiency

```
##        female:ID
## Down           0
## NotSig     15060
## Up             0
```

##### Sex/batch specific responses to choline supplementation

```
##        female:choline
## Down              446
## NotSig          14064
## Up                550
```

| | SYMBOL | FDR | logFC | DESCRIPTION |
|---|---|---|---|---|
| 15745 | LOC100360841 | 1.00e-07 | 1.6696958 | ribosomal protein L37 |
| 21872 | Giot1 | 2.00e-07 | 1.4838244 | gonadotropin inducible ovarian transcription factor 1 |
| 3006 | Ccdc82 | 5.00e-07 | -0.9075426 | coiled-coil domain containing 82 |
| 9417 | LOC685273 | 9.00e-07 | -1.3729108 | similar to U1 small nuclear ribonucleoprotein C (U1 snRNP protein C) (U1C protein) (U1-C) |
| 16696 | AC135409 | 9.00e-07 | -1.8740545 | NULL |
| 19561 | Capns1 | 1.50e-06 | 4.6802629 | calpain small subunit 1-like |
| 19711 | AABR07044509 | 3.40e-06 | -1.3539613 | NULL |
| 28245 | AABR07000733 | 5.30e-06 | 4.7380420 | NULL |
| 5580 | Scube1 | 1.21e-05 | -1.1604288 | signal peptide, CUB domain and EGF like domain containing 1 |
| 3780 | Fbln2 | 1.55e-05 | -1.1863910 | fibulin 2 |

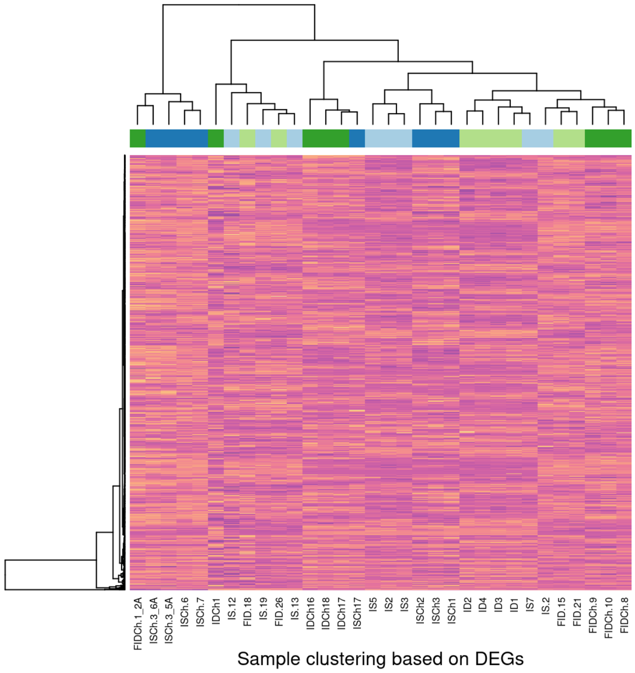


Sample clustering based on DEGs

```
## png
##   2
```

```
## Joining, by = c("sampleID", "sex", "iron", "choline", "diet")
```

A violin plot of the top 6 genes with the lowest FDR value in the sex:choline test (below) evidences different effects of choline supplementation on the expression profiles depending on the sex of the pup. For comparison, a similar plot using the expression values of 6 randomly-selected genes is also shown.

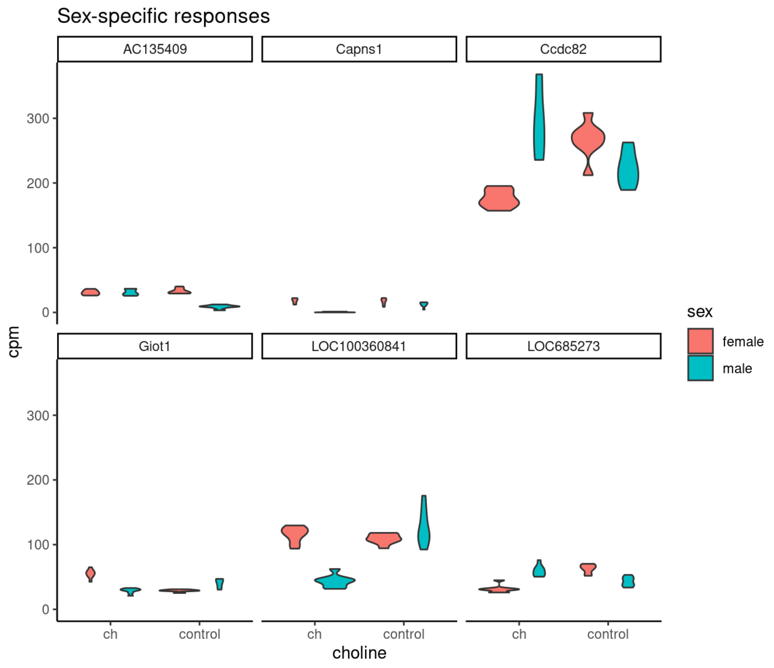


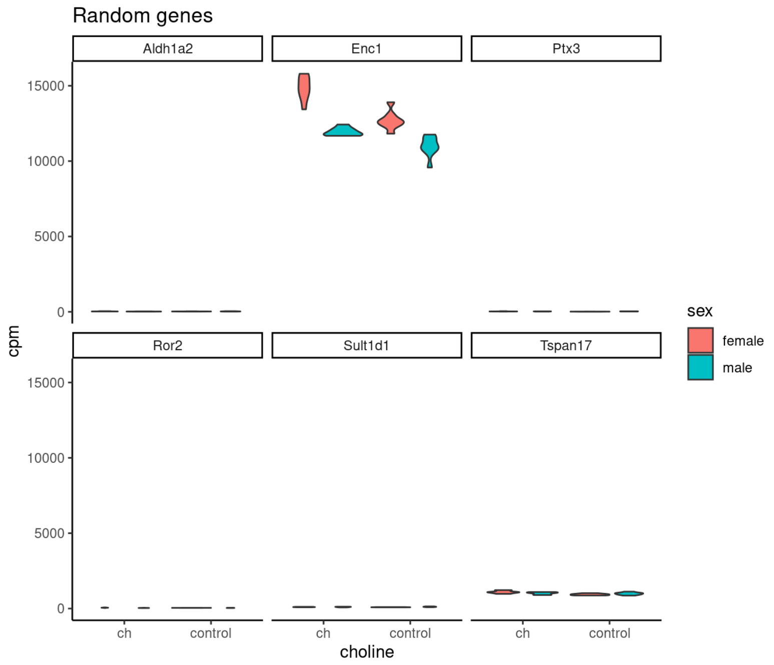


##### Iron:choline interaction

```
##        ID:choline
## Down            0
## NotSig      15060
## Up              0
```

### Analysis of Female Pups

Data from 16 female pups (4 from each diet combination) were used for the analysis. From the 32883 annotated genes, those shorter than 300 bp or with no expression in any of the dietary groups were filtered out, leaving a total of 15304 loci.

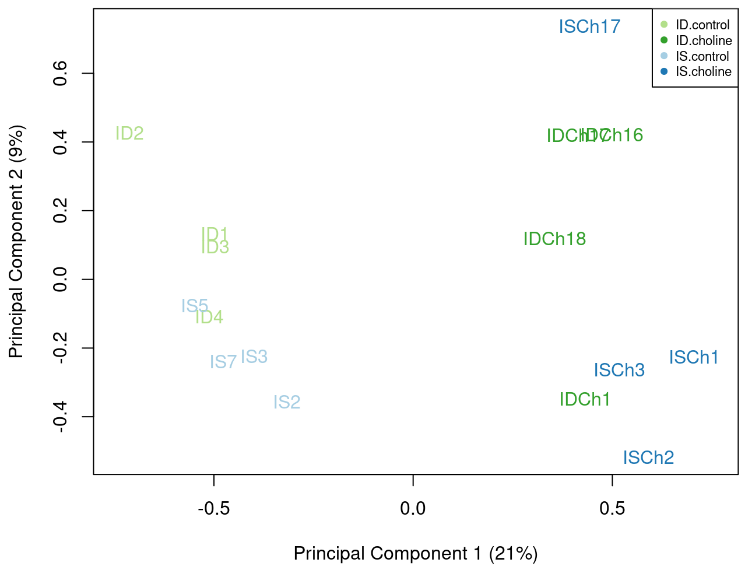


```
## png
##   2
```

The exploratory MDS plot shows that choline supplementation explains approximately 20% of the variability in the expression profiles of all female samples. Iron status does not separate samples as well as choline status.

#### Analysis of variance

```
## Saving 7 x 6 in image
```

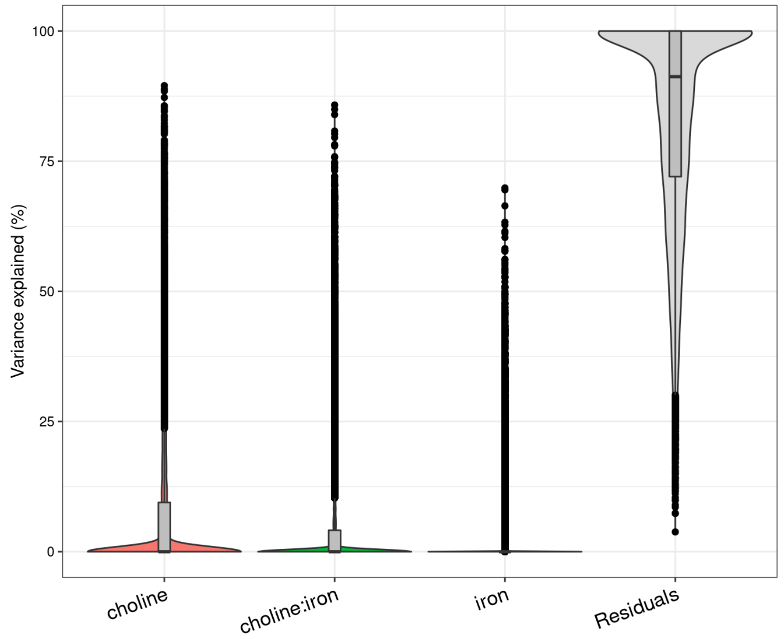


The top 100 genes that deviate from the genome-wide trend (outliers from the variance plot above) are selected and stored in the vp.txt files, the variance partition of the top 10 genes for each variable are plotted and saved as vp.pdf files.

```
## Saving 7 x 6 in image
## Saving 7 x 6 in image
## Saving 7 x 6 in image
```

#### Differential gene expression

To estimate the effect of dietary iron levels and choline supplementation, the expression profiles of each gene (y) was modeled as y = f(diet). The threshold for differential expression is defined as FDR < 0.05.

##### Effect of iron deficiency

- All ID vs all IS libraries

```
##        -1*IS.control -1*IS.choline 1*ID.control 1*ID.choline
## Down                                                        2
## NotSig                                                  14962
## Up                                                          1
```

- ID.control vs IS.control

```
##        -1*IS.control 1*ID.control
## Down                            3
## NotSig                      14958
## Up                              4
```

##### Effect of iron deficiency and choline supplementation

- ID.choline vs IS.choline

```
##        -1*IS.choline 1*ID.choline
## Down                            3
## NotSig                      14960
## Up                              2
```

Few genes were found as DE due to iron status in female pups. Regardless of choline supplementation status, 3 genes were differentially expressed in female pups with any ID diet compared to pups with any IS diet. Only 6 genes were DE between the ID.control vs IS.control diets and there were only 5 DEGs between the ID.choline and IS.choline groups. These results imply **no considerable difference between groups that can be explained by their dietary iron levels.**

##### Effect of choline supplementation

- All choline vs all control libraries

```
##        -1*IS.control 1*IS.choline -1*ID.control 1*ID.choline
## Down                                                      629
## NotSig                                                  13625
## Up                                                        711
```

| | SYMBOL | FDR | logFC | DESCRIPTION |
|---|---|---|---|---|
| 7382 | Dsp | 0.0e+00 | 5.822200 | desmoplakin |
| 8809 | Ift140 | 0.0e+00 | 1.436667 | intraflagellar transport 140 |
| 9254 | Ephx2 | 1.0e-07 | 7.233976 | epoxide hydrolase 2 |
| 13113 | Pdyn | 2.0e-07 | 3.117733 | prodynorphin |
| 2097 | Pld5 | 1.2e-06 | 3.774958 | phospholipase D family, member 5 |
| 9321 | AABR07001512 | 2.3e-06 | -6.380645 | NULL |
| 9412 | Bphl | 2.3e-06 | 1.959965 | biphenyl hydrolase like |
| 14369 | AABR07054368 | 2.3e-06 | -1.120208 | NULL |
| 9541 | Nqo2 | 2.3e-06 | -3.339293 | N-ribosyldihydronicotinamide:quinone reductase 2 |
| 210 | Snrpc | 2.3e-06 | 1.409270 | small nuclear ribonucleoprotein polypeptide C |

Choline supplementation did show a clear effect on the expression profiles of 1358 genes. Female pups with choline-supplemented diets showed 665 genes downregulated and 693 upregulated compared to non-supplemented diets, regardless of their dietary iron levels.

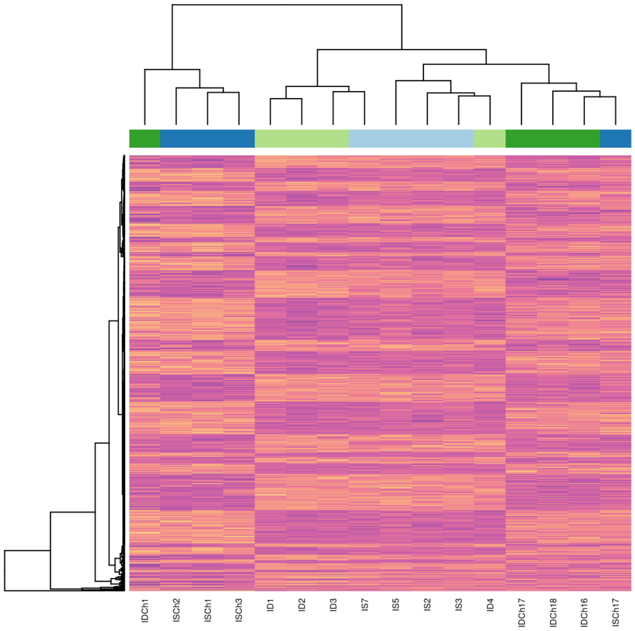


Sample clustering based on DEGs

```
## png
##   2
```

- IS.choline vs IS.control

```
##        -1*IS.control 1*IS.choline
## Down                          153
## NotSig                      14569
## Up                            243
```

| | SYMBOL | FDR | logFC | DESCRIPTION |
|---|---|---|---|---|
| 7382 | Dsp | 0.0000005 | 2.8703161 | desmoplakin |
| 21877 | AABR07031918 | 0.0000176 | 4.9718128 | small conductance calcium-activated potassium channel protein 2-like |
| 9254 | Ephx2 | 0.0000293 | 3.4166906 | epoxide hydrolase 2 |
| 14369 | AABR07054368 | 0.0000371 | -0.7025595 | NULL |
| 6582 | Cemip | 0.0000392 | 0.5359933 | cell migration inducing hyaluronidase |
| 8809 | Ift140 | 0.0000410 | 0.5476669 | intraflagellar transport 140 |
| 13113 | Pdyn | 0.0000410 | 1.4897360 | prodynorphin |
| 2097 | Pld5 | 0.0000437 | 2.0447282 | phospholipase D family, member 5 |
| 21872 | Giot1 | 0.0002549 | 1.0452742 | gonadotropin inducible ovarian transcription factor 1 |
| 556 | Slc29a4 | 0.0002549 | 0.8779574 | solute carrier family 29 member 4 |

Only 390 DEGs were found in the IS.choline vs IS.control comparison. However, sample clustering based on this set of genes does follow the sample choline status.

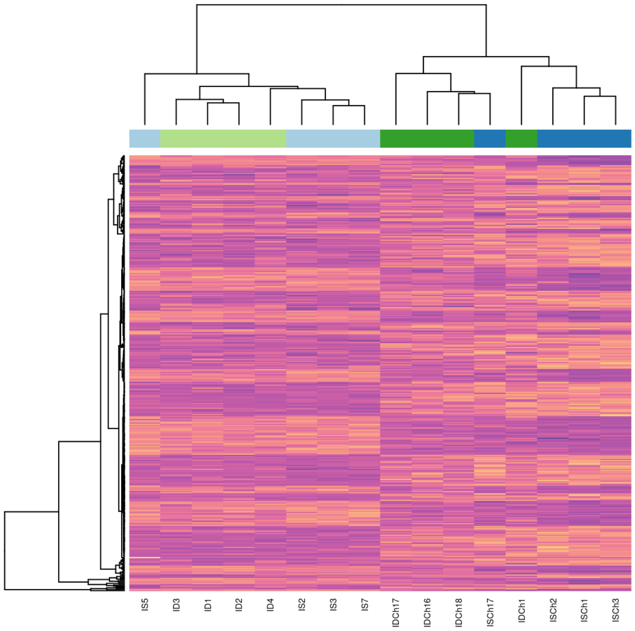


Sample clustering based on DEGs

```
## png
##   2
```

##### Rescued effects of choline supplementation

- ID.choline vs. IS.control

```
##        -1*IS.control 1*ID.choline
## Down                          111
## NotSig                      14722
## Up                            132
```

| | SYMBOL | FDR | logFC | DESCRIPTION |
|---|---|---|---|---|
| 8809 | Ift140 | 0.0000002 | 0.8748123 | intraflagellar transport 140 |
| 7382 | Dsp | 0.0000003 | 2.8111969 | desmoplakin |
| 9254 | Ephx2 | 0.0000042 | 3.8584168 | epoxide hydrolase 2 |
| 9321 | AABR07001512 | 0.0000088 | -4.9659677 | NULL |
| 13113 | Pdyn | 0.0000280 | 1.5622479 | prodynorphin |
| 9541 | Nqo2 | 0.0000433 | -2.0112891 | N-ribosyldihydronicotinamide:quinone reductase 2 |
| 9412 | Bphl | 0.0000921 | 1.0863142 | biphenyl hydrolase like |
| 2097 | Pld5 | 0.0001523 | 1.8661246 | phospholipase D family, member 5 |
| 3780 | Fbln2 | 0.0001854 | -0.8555052 | fibulin 2 |
| 210 | Snrpc | 0.0004140 | 0.6875763 | small nuclear ribonucleoprotein polypeptide C |

Female pups with an ID.choline diet were compared to female pups with and IS.control diet. A total of 242 genes were differentially expressed between groups. The 10 genes with lowest FDR values are shown.

The heatmap below shows that most of the difference among expression values in this set of genes can be explained from the choline supplementation (i.e. the expression values of most DEGs are similar in all choline diets and different from all control diets) rather than the difference in iron status.

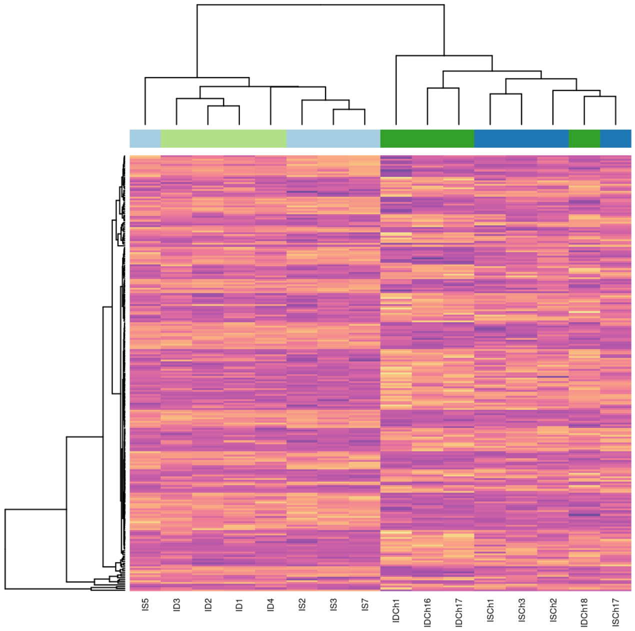


Sample clustering based on DEGs

```
## png
##   2
```

##### Additional tests

The following tests aim to understand the different effects of choline supplementation in pups with an ID and the possible interaction between iron status and choline supplementation.

- Effect of choline in ID diets = ID.choline vs ID.control
- Iron:choline interaction = Effect of choline in ID diets - Effect of choline in IS diets

**Results:**

- Effect of choline in ID diets

```
##        -1*ID.control 1*ID.choline
## Down                          312
## NotSig                      14396
## Up                            257
```

| | SYMBOL | FDR | logFC | DESCRIPTION |
|---|---|---|---|---|
| 8809 | Ift140 | 1.00e-07 | 0.8889999 | intraflagellar transport 140 |
| 7382 | Dsp | 2.00e-07 | 2.9518839 | desmoplakin |
| 9254 | Ephx2 | 4.00e-06 | 3.8172856 | epoxide hydrolase 2 |
| 9321 | AABR07001512 | 4.00e-06 | -5.1810715 | NULL |
| 13113 | Pdyn | 1.63e-05 | 1.6279966 | prodynorphin |
| 3006 | Ccdc82 | 5.19e-05 | -0.7708208 | coiled-coil domain containing 82 |
| 4892 | Sez6 | 6.78e-05 | -0.4034003 | seizure related 6 homolog |
| 210 | Snrpc | 7.47e-05 | 0.7950335 | small nuclear ribonucleoprotein polypeptide C |
| 2444 | Rnd3 | 7.47e-05 | 0.6851752 | Rho family GTPase 3 |
| 9412 | Bphl | 8.10e-05 | 1.0712049 | biphenyl hydrolase like |

Female pups with ID and choline-supplemented diets had 309 genes downregulated and 244 upregulated when compared to those with ID.control diets. The top 10 genes with the lowest FDR values from this comparison are displayed.

Using only the DEGs to cluster all libraries (heatmap below) shows that samples clearly separate by choline supplementation status. Within the control group (no choline), samples also separate depending on their iron status, but within the choline group, the sample clustering does not follow iron status. This implies that **choline supplementation modifies the expression profiles of this set of genes in female pups with ID diets so that they are more similar to the expression profiles of pups with IS diets.**

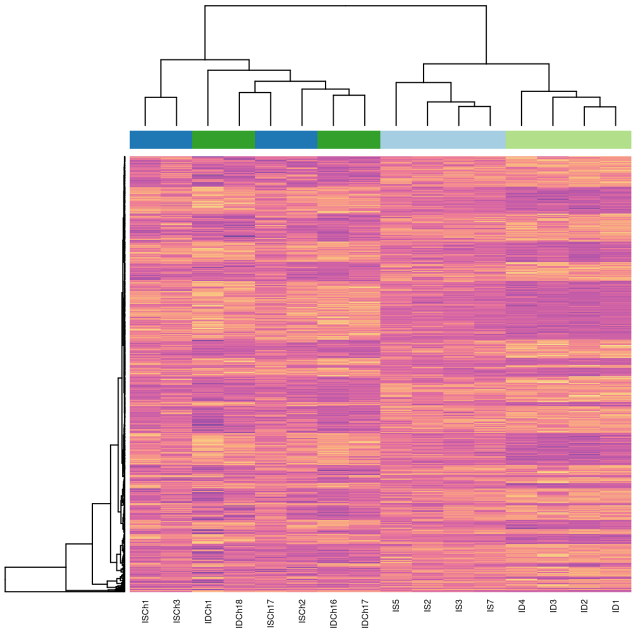


Sample clustering based on DEGs

```
## png
##   2
```

- Iron:choline interaction

```
##        1*IS.control -1*IS.choline -1*ID.control 1*ID.choline
## Down                                                       1
## NotSig                                                 14964
## Up                                                         0
```

**The IS.choline vs IS.control and ID.choline vs ID.control tests evidence that choline supplementation affects a different set of genes, depending on the dietary iron levels.** However, the interaction between choline and iron status (choline supplementation affecting a specific gene "x" differently depending on the dietary iron levels) was tested and found only found 1 DEG. This suggests that there is no need for an interaction term to explain the observed data (i.e. **there is no evidence of a different effect of choline on a specific gene at different dietary iron levels**).

#### Analysis of Male Pups

Data from 16 male pups (4 from each diet combination) were used for the analysis. From the 32883 genes detected in the study, those shorter than 300 bp or with no expression in any of the dietary groups were filtered out, leaving a total of 14585 loci.

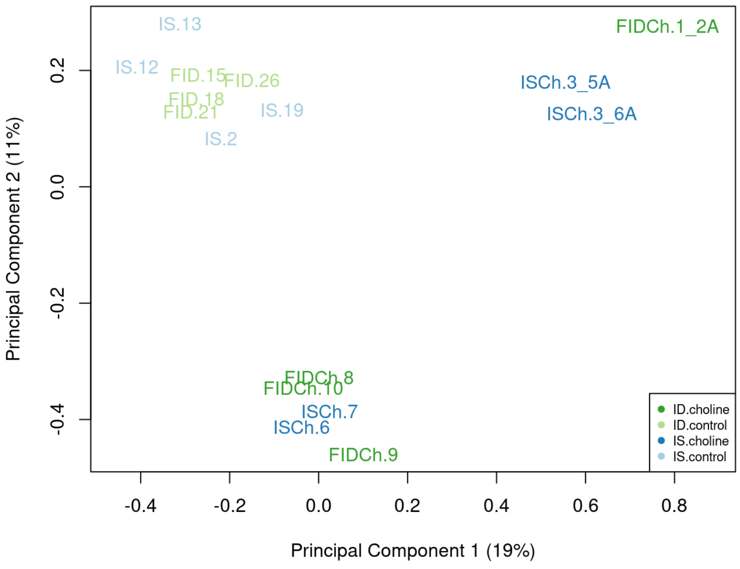

```
## png
##   2
```

The exploratory MDS plot shows that the principal sources of variation among samples do not correspond clearly with any of the explanatory variables of the study (iron and choline). However, choline supplementation seems to have a stronger effect on the expression profiles than the iron levels since all control samples cluster together while choline-supplemented diets show greater heterogeneity in their expression profiles.

#### Analysis of variance

```
## Saving 7 x 6 in image
```

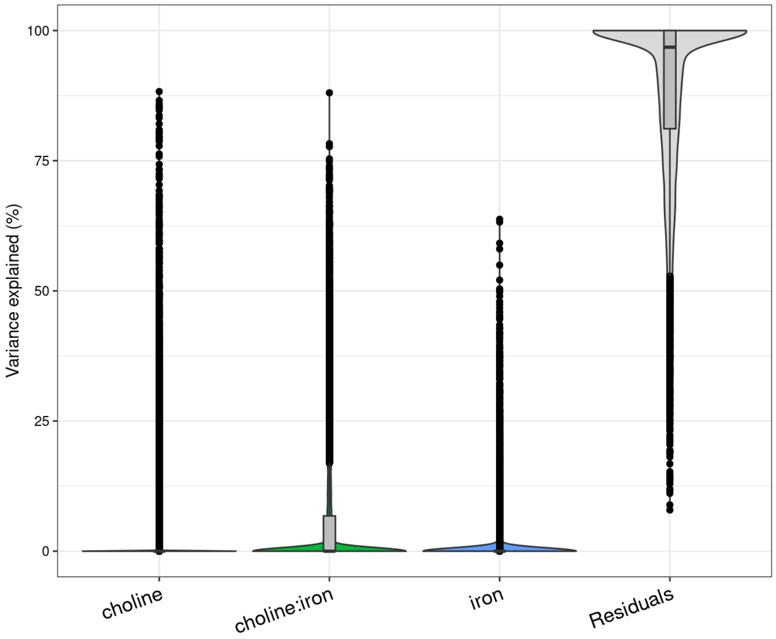


The top 100 genes that deviate from the genome-wide trend (outliers from the variance plot above) are selected and stored in the vp.txt files, the variance partition of the top 10 genes for each variable are plotted and saved as vp.pdf files.

```
## Saving 7 x 6 in image
## Saving 7 x 6 in image
## Saving 7 x 6 in image
```

#### Differential gene expression

To estimate the effect of dietary iron levels and choline supplementation, all 16 male pup libraries were used in a full comparison. The expression profile (y) was modeled as y = f(diet).

##### Effect of iron deficiency

- All ID vs all IS libraries

```
##        -1*IS.control -1*IS.choline 1*ID.control 1*ID.choline
## Down                                                       0
## NotSig                                                 14305
## Up                                                         0
```

- ID.control vs IS.control

```
##        -1*IS.control 1*ID.control
## Down                            0
## NotSig                      14305
## Up                              0
```

There are no DEGs between ID and IS diets in male pups.

##### Effect of iron deficiency and choline supplementation

- ID.choline vs IS.choline

```
##        -1*IS.choline 1*ID.choline
## Down                            0
## NotSig                      14305
## Up                              0
```

There are no DEGs between ID.choline and IS.choline diets in male pups.

##### Effect of choline supplementation

- All choline vs all control libraries

```
##        -1*IS.control 1*IS.choline -1*ID.control 1*ID.choline
## Down                                                      77
## NotSig                                                 14173
## Up                                                        55
```

| | SYMBOL | FDR | logFC | DESCRIPTION |
|---|---|---|---|---|
| 5197 | LOC680121 | 4.80e-06 | 1.3593262 | similar to heat shock protein 8 |
| 15745 | LOC100360841 | 5.70e-06 | -3.1753314 | ribosomal protein L37 |
| 13994 | AABR07072078 | 6.30e-06 | -2.9027082 | NULL |
| 15465 | Mt | 6.40e-06 | 0.9368608 | mitochondrially encoded ATP synthase 8 |
| 18038 | LOC100365810 | 1.27e-05 | -2.4192601 | 40S ribosomal protein S17-like |
| 19711 | AABR07044509 | 1.27e-05 | 2.7779409 | NULL |
| 15366 | AABR07025089 | 1.34e-05 | -4.0638918 | NULL |
| 26675 | AC096600 | 1.34e-05 | 1.8897057 | NULL |
| 15739 | Cox6b1 | 1.48e-05 | 1.1244770 | cytochrome c oxidase subunit 6B1 |
| 16696 | AC135409 | 1.52e-05 | 3.5557405 | NULL |

Choline supplementation significantly modified the expression profiles of 147 genes.

Using this set of genes to cluster the samples does not show the expected clasification based on dietary group.

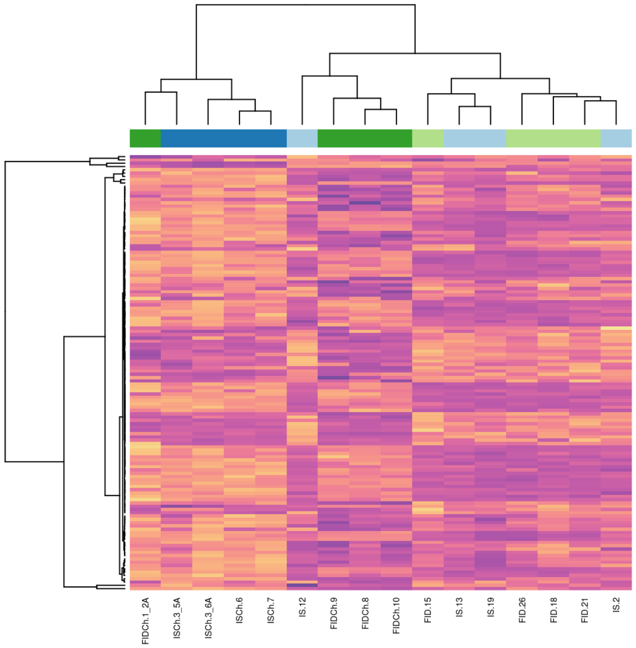


Sample clustering based on DEGs

```
## png
##   2
```

- IS.choline vs IS.control

```
##        -1*IS.control 1*IS.choline
## Down                           21
## NotSig                      14268
## Up                             16
```

| | SYMBOL | FDR | logFC | DESCRIPTION |
|---|---|---|---|---|
| 5197 | LOC680121 | 0.0002304 | 0.7495607 | similar to heat shock protein 8 |
| 15745 | LOC100360841 | 0.0002304 | -1.7422494 | ribosomal protein L37 |
| 13994 | AABR07072078 | 0.0003313 | -1.5854002 | NULL |
| 19711 | AABR07044509 | 0.0013162 | 1.4598201 | NULL |
| 18038 | LOC100365810 | 0.0013162 | -1.2489269 | 40S ribosomal protein S17-like |
| 25725 | LOC100363531 | 0.0015710 | -0.6067593 | LRRGT00182-like |

| | SYMBOL | FDR | logFC | DESCRIPTION |
|---|---|---|---|---|
| 27759 | AC128792 | 0.0015710 | -1.0074914 | NULL |
| 14717 | Mt | 0.0015710 | -0.6801499 | mitochondrially encoded NADH dehydrogenase 2 |
| 14591 | Mt | 0.0026441 | -0.5586373 | mitochondrially encoded NADH dehydrogenase 1 |
| 19561 | Capns1 | 0.0026441 | -3.9447359 | calpain small subunit 1-like |

Only 49 DEGs were found in the IS.choline vs IS.control comparison. Sample clustering based on this set of genes is shown below and resembles the expected classification based on choline status. Interestingly, the different loci for the gene Mt have different responses depending on the dietary group.

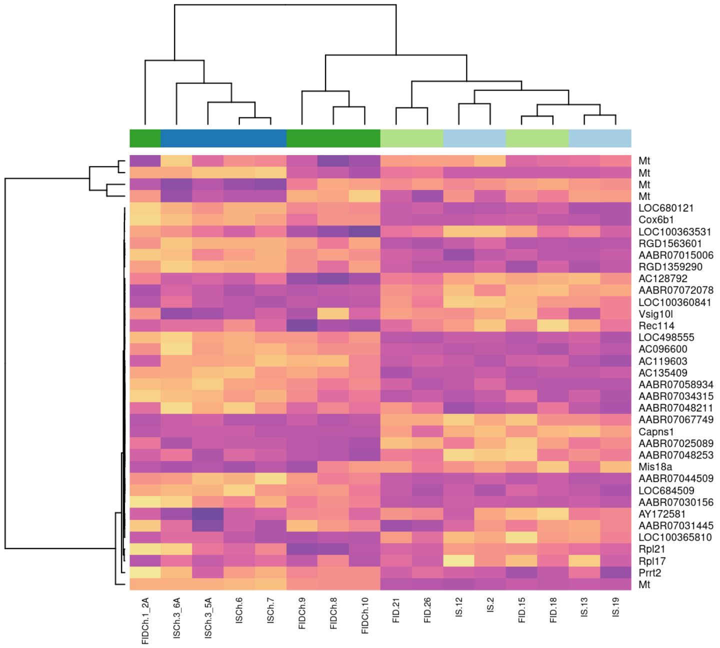

Sample clustering based on DEGs

```
## png
##   2
```

##### Rescued effects of choline supplementation

- ID.choline vs. IS.control

```
##        -1*IS.control 1*ID.choline
## Down                            34
## NotSig                       14251
## Up                              20
```

| | SYMBOL | FDR | logFC | DESCRIPTION |
|---|---|---|---|---|
| 5197 | LOC680121 | 0.0000433 | 0.8609070 | similar to heat shock protein 8 |
| 15745 | LOC100360841 | 0.0005322 | -1.7201220 | ribosomal protein L37 |
| 27759 | AC128792 | 0.0010649 | -1.1809582 | NULL |
| 18038 | LOC100365810 | 0.0011604 | -1.2913586 | 40S ribosomal protein S17-like |
| 13994 | AABR07072078 | 0.0011604 | -1.4150718 | NULL |
| 15465 | Mt | 0.0011604 | 0.4581442 | mitochondrially encoded ATP synthase 8 |
| 19561 | Capns1 | 0.0011604 | -6.6265691 | calpain small subunit 1-like |
| 11832 | AABR07015006 | 0.0012625 | 0.7066175 | similar to Heterous nuclear ribonucleoprotein A1 (Helix-destabilizing protein) (Single-strand RNA-binding protein) (hnRNP core protein A1) (HDP) |
| 25725 | LOC100363531 | 0.0013069 | -0.6235961 | LRRGT00182-like |
| 15739 | Cox6b1 | 0.0013442 | 0.5939556 | cytochrome c oxidase subunit 6B1 |

Male pups with an ID.choline diet were compared to the male pups with and IS.control diet. A total of 64 genes were differentially expressed between groups. Pups with ID.choline diets had 42 genes downregulated and 22 upregulated with respect to the IS.control. The 10 genes with lowest FDR values are shown.

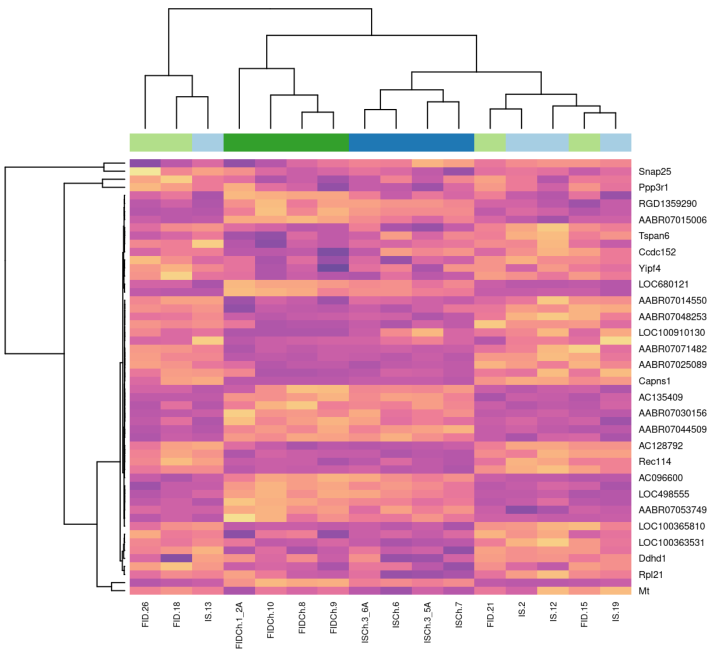


Sample clustering based on DEGs

```
## png
##   2
```

When using the expression values of these genes to cluster all samples, it becomes clear that most of the DEGs found correlate with the choline supplementation i.e. the expression values of most DEGs are similar in all choline diets and different from all control diets regardless of the iron status, except, perhaps, the gene cluster to which Tspan6, Yipf4 and Ccdc152 belong.

##### Additional tests

The following tests aim to understand the different effects of choline supplementation in pups with either an ID or an IS diet and the possible interaction between iron status and choline supplementation.

- Effect of choline in ID diets = ID.choline vs ID.control
- Iron:choline interaction = Effect of choline in ID diets - Effect of choline in IS diets

**Results:**

- Effect of choline in ID diets

```
##        -1*ID.control 1*ID.choline
## Down                          190
## NotSig                      14029
## Up                             86
```

| | SYMBOL | FDR | logFC | DESCRIPTION |
|---|---|---|---|---|
| 15465 | Mt | 0.0007709 | 0.5352957 | mitochondrially encoded ATP synthase 8 |
| 14950 | RGD1563601 | 0.0011144 | 0.9152101 | similar to Tpi1 protein |
| 15366 | AABR07025089 | 0.0011144 | -2.2343778 | NULL |
| 16696 | AC135409 | 0.0011144 | 2.0844655 | NULL |
| 5197 | LOC680121 | 0.0011144 | 0.6097655 | similar to heat shock protein 8 |
| 15739 | Cox6b1 | 0.0015261 | 0.6022851 | cytochrome c oxidase subunit 6B1 |
| 26675 | AC096600 | 0.0016620 | 0.9849016 | NULL |
| 16516 | LOC498555 | 0.0016620 | 0.9015465 | similar to 60S acidic ribosomal protein P2 |
| 15745 | LOC100360841 | 0.0017332 | -1.4330821 | ribosomal protein L37 |
| 32205 | LOC680491 | 0.0017332 | -0.6613195 | similar to heat shock protein 1, alpha |

Male pups with ID and choline-supplemented diets had 181 genes downregulated and 104 upregulated when compared to those with ID.control diets. The top 10 genes with the lowest FDR values from this comparison are displayed.

Using only these genes to cluster all male libraries (heatmap below), samples do not show a clear separation by choline supplementation and iron status in the contraol samples as it did for the female samples.

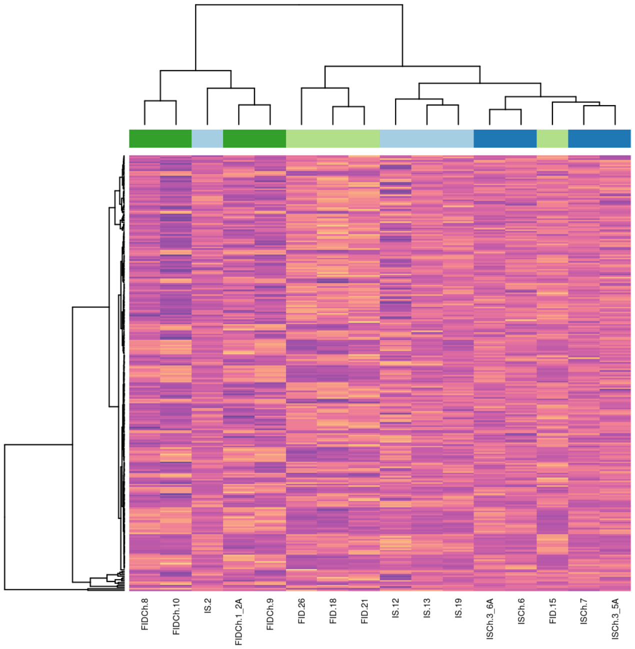

Sample clustering based on DEGs

```
## png
##   2
```

- Iron:choline interaction

```
##        1*IS.control -1*IS.choline -1*ID.control 1*ID.choline
## Down                                                       1
## NotSig                                                 14304
## Up                                                         0
```

Choline supplementation affects a different set of genes depending on the dietary iron levels. However, the interaction between choline and iron status only found 1 DEG. This suggests that choline supplementation does not affect a specific gene "x" differently depending on the dietary iron levels.

### Conclusions

A joint analysis of all RNA-seq libraries can not support any conclusion related to the differential effects in female vs male rats given the different batches and technologies used for the growth and sequencing of each sex. However, an exploratory analysis of the variance and differential gene expression in the joint datasets suggested the existence of interaction effects between sex/batch and the study variables, i.e. sex/batch-specific effects to choline suplemmentation or iron status. To better understand this, each sex/batch was analyzed separately and the trends found *within* each one are compared with the other.

#### Analysis of variance

The genome wide expression variance among samples do not correlate with the explanatory variables of the study. This is, the variance in gene expression for most genes is due to other factors that can not be controlled in the study such as individual genetic variation.

In the joint dataset, the variation across sex/batch/technology explains a median of 14.6 % of the variation in gene expression, while the median effects of choline, iron and their interactions are negligible after correcting for all the others. Nontheless, such analysis allows finding genes that do not follow the genome-wide trend, i.e. genes with expression profiles that respond to choline, iron, sex and their interactions, even if they do not cross the threshold for differential gene expression. The top 100 genes with the highest percent of variance explained by each of the study variables were found (.vp.txt files) and the variance partition of the top 10 plotted as barplots (.vp.pdf files).

#### Differential gene expression

Summary of the joint analysis:

| variable | comparison | DEGs |
|---|---|---|
| Iron | all ID vs all IS | 0 |
| Choline | all choline vs all control | 101 |
| Interaction | sex:iron | 0 |
| Interaction | sex:choline | 996 |
| Interaction | iron:choline | 0 |

A summary table with the DGE results comparing the datasets separately by batch/sex is presented below:

| variable | comparison | female.DEGs | male.DEGs |
|---|---|---|---|
| Iron | all ID vs all IS | 3 | 0 |
| Iron | ID.control vs IS.control | 7 | 0 |

| variable | comparison | female.DEGs | male.DEGs |
|---|---|---|---|
| Iron | ID.choline vs IS.choline | 5 | 0 |
| Choline | all choline vs all control | 1340 | 132 |
| Choline | ID.choline vs ID.control | 569 | 276 |
| Choline | IS.choline vs IS.control | 396 | 37 |
| Iron:choline | Interaction | 1 | 1 |
| Rescued effects | ID.choline vs IS.control | 243 | 54 |

**DEGs related to iron deficiency**

There is no evidence of a considerable difference between the hippocampal transcriptomes of pups with IS and ID diets, regardless of their sex or choline supplementation status.

**DEGs related to choline supplementation**

Choline supplementation was the explanatory variable with the most pronounced effect in the transcriptomic differences for both sexes. About 20 % of the variability in the transcriptomes of female pups can be explained by the samples' choline supplementation status as observed in the MDS plot of female samples. For males, the effects of choline supplementation are less uniform than for females, i.e. choline-supplemented diets have different transcriptomic profiles that do not correlate with any known variable.

Choline supplementation did not affect a specific gene differently depending on the dietary iron levels for any sex (i.e. the interaction term iron:choline was not significant).

Separate analyses between IS (IS.choline vs IS.control) or ID diets (ID.choline vs ID.control) showed that ID diets correlate with a more pronounced effect from choline supplementation than IS diets. Specifically, there are more DEGs in the ID.choline vs ID.control comparison than in the IS.choline vs IS.control comparison for both sexes.

**DEGs related to the rescued effects of choline supplementation**

The researcher defined "rescued effects of choline supplementation" as the comparison ID.choline vs IS.control. Choline supplementation status can explain most of the DEGs found with this comparison, i.e. genes upregulated in the ID.choline group compared to the IS.control group are also upregulated in the IS.choline group compared to the IS.control. However, a few genes do not follow this trend.

### GSEA and ORA notes

Command used:

```
run_clusterProfiler.R -f females.Ch.deg.txt -s Rattus_norvegicus -d females/choline -i SYMBOL -p 0.1 -n 15 -b ALL --gsea TRUE --prerankMethod pval
```

ORA options: It only uses DEGs, since the number of DEGs is not big enough for most of the comparisons (hsould be between 200 - 400 aproximately), it is not useful except for the choline tests. - FDR cutoff for DE: 0.1 - Reports the top 15 hits - Uses the 200 genes with lowest FDR as long as FDR is < 0.1 - Databases: all GO branches, KEGG and REACTOME

GSEA options: Uses a ranked list regardless of differential expression. This is the recommended way to proceed for further comparisons. All lists of DE tests and variance analysis were used for the GSEA. - Ranked by -log10(PVal)*sign(FC) for DE lists and by %variance explained for variance lists. - Database: H - For ranked variance lists: genes with same mapping to the human ortholog were collapsed and the tests used 1000 permutations.
